# Mitochondrial Phosphopantetheinylation is Required for Oxidative Function

**DOI:** 10.1101/2024.05.09.592977

**Authors:** Pieter R. Norden, Riley J. Wedan, Samuel E. J. Preston, Morgan Canfield, Naomi Graber, Jacob Z. Longenecker, Olivia A. Pentecost, Elizabeth McLaughlin, Madeleine L. Hart, Sara M. Nowinski

## Abstract

4’-phosphopantetheinyl (4’PP) groups are essential co-factors added to target proteins by <u>p</u>hospho<u>p</u>antetheinyl transferase (PPTase) enzymes. Although mitochondrial 4’PP-modified proteins have been described for decades, a mitochondrially-localized PPTase has never been found in mammals. We discovered that the cytoplasmic PPTase <u>a</u>mino<u>a</u>dipate <u>s</u>emialdehyde <u>d</u>ehydrogenase <u>p</u>hospho<u>p</u>antetheinyl transferase (AASDHPPT) is required for mitochondrial respiration and oxidative metabolism. Loss of AASDHPPT results in failed 4’PP modification of the mitochondrial acyl carrier protein and blunted activity of the mitochondrial fatty acid synthesis (mtFAS) pathway. We found that in addition to its cytoplasmic localization, AASDHPPT localizes to the mitochondrial matrix via an N-terminal mitochondrial targeting sequence contained within the first 20 amino acids of the protein. Our data show that this novel mitochondrial localization of AASDHPPT is required to support mtFAS activity and oxidative function. We further identify five variants of uncertain significance in *AASDHPPT* that are likely pathogenic in humans due to loss of mtFAS activity.

## Main

4’-phosphopantentheinylation is an essential post translational modification (PTM) whereby a 4’-phosphopantetheine (4’PP) group, derived from coenzyme A (CoA), is covalently attached to target proteins by a phosphopantetheinyltransferase (PPTase) enzyme^1^. A total of six proteins in the mammalian proteome have been found to have a 4’PP-modified serine: three cytoplasmic proteins (Fatty Acid Synthase, FASN; Aminoadipate semialdehyde dehydrogenase, AASDH; and Aldehyde Dehydrogenase 1 Family Member L1, ALDH1L1), and three mitochondrial proteins (NADH:Ubiquinone Oxidoreductase Subunit AB1, NDUFAB1; Aldehyde Dehydrogenase 1 Family Member L2, ALDH1L2; and mitochondrial Dehydrogenase/Reductase 2, DHRS2)^1^. In mammalian genomes, only one PPTase has been annotated—<u>a</u>mino<u>a</u>dipate-<u>s</u>emialdehyde <u>d</u>e<u>h</u>ydrogenase-<u>p</u>hospho<u>p</u>antetheinyl transferase (AASDHPPT)—which was thought to exist solely in the cytoplasm^2^. This contrasts with other species such as *Arabidopsis thaliana* and *Saccharomyces cerevisiae,* which have dedicated mitochondrial PPTases, the activity of which cannot be compensated by cytoplasmic PPTase proteins^3,4^. It was therefore unclear how 4’PP is attached to mitochondrial proteins in mammalian cells.

Since no mammalian mitochondrial PPTase had ever been described, we questioned whether an unidentified PPTase existed in mammals, or conversely if AASDHPPT (hereafter abbreviated PPT) was important for mitochondrial function regardless of its reported cytoplasmic localization. One prior study found that PPT could add a 4’PP group to purified mitochondrial proteins *in vitro*, but concluded that its activity was greater than 90% cytoplasmic, raising questions as to whether this activity was biologically relevant^2^. We therefore set out to understand whether PPT is required for mitochondrial function and, if so, how it modifies mitochondrial targets despite existing in separate cellular compartments.

To test whether PPT is required for mitochondrial oxidative function, we used a CRISPR/Cas-9-based strategy to mutate *Aasdhppt* in C2C12 mouse skeletal muscle myoblasts. Complete loss of PPT was incompatible with growth; however, we generated two clonal cell lines with hypomorphic mutations in *Aasdhppt* (Extended Data Fig. 1A) that resulted in decreased PPT protein levels (PPT-1 and PPT-2, Fig. 1A). These results were confirmed by siRNA-mediated knockdown of the same target (siPPT-1 and siPPT-2, Fig. 1A). Of note, C2C12 cells have been shown to be near-tetraploid^5,6^. Consequently, we found three alleles with unique PPT gene sequences in each of our PPT-deficient C2C12 cell lines (Extended

**Figure 1.**
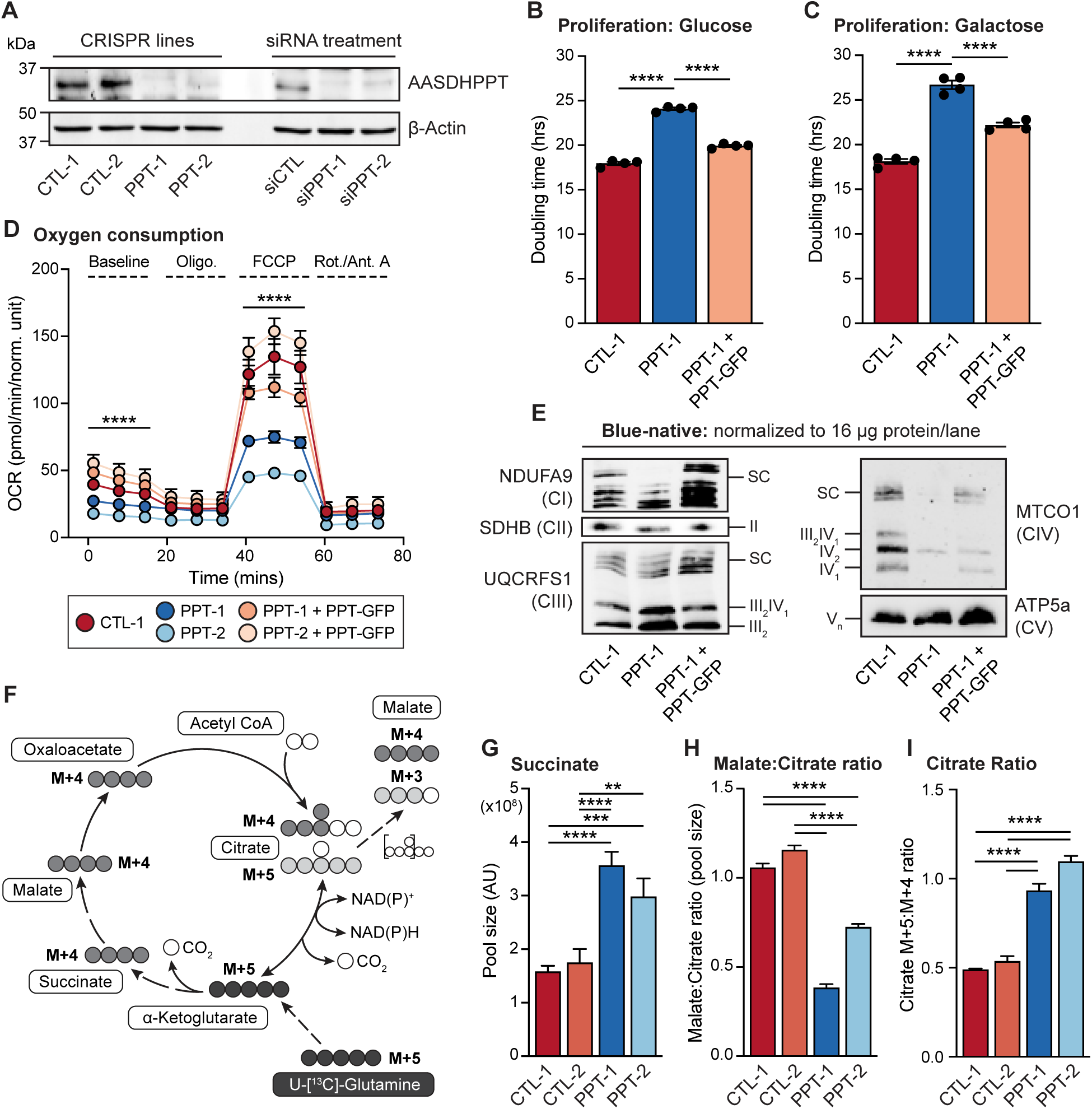
AASDHPPT is required for mitochondrial oxidative metabolism. (A) Whole cell lysates were separated by SDS-PAGE and immunoblotted for the denoted targets in clonal control, *Aasdhppt*-deficient, and siRNA-transfected cells. Data are representative of 3 biological replicates. (B-C) Cells were grown in either 25 mM glucose (B) or 10 mM galactose (C) in an IncuCyte® system with images captured every 3 hours for 4 days or 2 days respectively. Data represent the mean doubling time calculated from technical replicates of growth curves; error bars denote SEM **** = p <0.0001. Data are representative of 1 experiment from n > 3 biological replicates. (D) Seahorse mitochondrial stress test for Oxygen Consumption Rate (OCR) of indicated cell lines expressing mtDSRed or PPT-GFP. Data are representative of 1 experiment from 3 biological replicates. Error bars represent SD. Significance determined at indicated time points using one-way ANOVA, **** = p <0.0001 (E) Blue-native PAGE separation of isolated mitochondrial protein complexes from denoted cell lines expressing mtDSRed or PPT-GFP, followed by immunoblot with the indicated antibodies. Data are representative of n > 3 biological replicates. (F) Schematic of isotopomeric labeling patterns upon substitution of U^13^C-glutamine for unlabeled glutamine. Open circles indicate ^12^C carbons, where circles with grey, red, or blue, indicate ^13^C carbons. (G-H) Technical triplicate samples (n=3) of the indicated cell lines cultured in U^13^C-glutamine for the indicated time points, harvested, and analyzed via LC-MS. Steady state metabolite pool sizes and ratios are shown for the 4-hour labeling time point. (I) Ratio of Citrate M+5 to Citrate M+4 at the four-hour time point. Error bars represent SEM (B, C), SD (D, G, H, I). ** = p<0.01, *** = p<0.001, **** = p<0.0001 as determined by ANOVA followed by Tukey’s multiple comparisons test.

Data Fig. 1A). In both cases, the top listed allele in Extended Data Fig. 1A was found about twice as frequently as the other two sequences in each cell line, likely indicating that there are two alleles with this sequence. Further, our sequencing analysis revealed that the PPT-2 cells still harbor at least one wild type allele, which corresponded to a low level of residual wild type PPT protein expression, also visible by western blot (Fig. 1A and Extended Data Fig. 1A). PPT-2 thus often displays a milder phenotype when compared directly with PPT-1, even though both exhibit a loss of PPT protein abundance (Fig. 1A).

We next asked whether PPT deficiency resulted in altered respiratory-dependent growth by culturing cells in media with either glucose or galactose, the latter requiring functional mitochondrial respiration. Both PPT-deficient cell lines showed significantly increased doubling times in both glucose-and galactose-containing media, which was rescued by transient re-expression of full length PPT with a C-terminal GFP-Flag tag (PPT-GFP) (Fig. 1B-C, Extended Data Fig. 1B-C). Next, we performed a Seahorse mitochondrial stress test and observed that PPT-deficient mutants exhibited a reduction in both basal respiration rate and maximal respiration rate compared to controls, rescued by re-expression of wild type PPT-GFP (Fig. 1D, Extended Data Fig. 1D). To assess whether PPT-deficient respiratory function was associated with perturbed OXPHOS complex expression and assembly, we performed blue-native PAGE on isolated mitochondria from PPT-deficient and control cells. PPT-deficient mutants displayed reduced levels of fully assembled electron transport chain (ETC) complex I (CI), complex II (CII) and complex IV (CIV). ATP synthase (CV) and complex III (CIII) were not affected by PPT-deficiency, but reductions in CIII-containing supercomplexes (SC) were observed. Lastly, assemblies of CI, II and IV were rescued by re-expression of wild type PPT-GFP (Fig. 1E, Extended Data Fig. 1E). Together, these data demonstrate that PPT deficiency results in dysfunctional mitochondrial respiratory and ETC-deficient phenotypes.

ETC impairments such as complex I dysfunction are well known to affect mitochondrial metabolism and alter tricarboxylic acid (TCA) cycle flux and nutrient utilization^7–9^. Since PPT-deficient cells display reduced levels of ETC complexes, we reasoned that they may also display altered TCA metabolism. To assess TCA cycle activity, we cultured cells in fully labeled ^13^C-glutamine (U-[^13^C]-Glutamine) for up to 6 hours and evaluated TCA cycle intermediates for relative abundance and carbon contribution from U-[^13^C]-labeled glutamine. This approach allowed us to assess reductive (light gray) versus oxidative (dark gray) metabolism of glutamine (Fig. 1F) both at steady state and over time. At steady state, PPT-deficient cells demonstrated an increase in the pool size of succinate and a decrease in downstream metabolites such as fumarate, malate, and aspartate (Fig. 1G, Extended Data Fig. 1F-H). Citrate and ⍺-ketoglutarate levels were variable (Extended Data Fig. 1I and J); however, the ratio of malate abundance to citrate abundance was consistently reduced in PPT mutants compared to controls (Fig. 1H). The reduction of TCA metabolites downstream of succinate relative to citrate suggests decreased oxidative TCA processing in PPT-deficient cells.

To assess whether this was indeed the case, we examined citrate labeling patterns from uniformly labeled glutamine. In general, M+4 citrate results from the oxidation of glutamine to oxaloacetate, whereas M+5 citrate results from the reductive carboxylation of glutamine to citrate. Increased reductive carboxylation is a common feature of respiratory-deficient cells^7,8^, because it affords cells an alternative pathway to regenerate NAD^+^. In agreement with our pool size data, which suggested a reduction in oxidative TCA processing, PPT-deficient cells showed reduced M+4 citrate and increased M+5 citrate compared to controls, indicating reduced oxidative TCA processing and increased reductive carboxylation of glutamine (Extended Data Fig. 1K). This is further supported by the increase in the ratio of citrate M+5:citrate M+4 at isotopic steady state (Fig. 1I). Accordingly, we also observe a decrease in downstream M+4 malate (Extended Data Fig. 1L), likely due to a reduction M+4 citrate. In sum, the glutamine labeling patterns in PPT-deficient cells demonstrate increased reductive carboxylation of glutamine, which is typical of cells with ETC impairments^5^. Collectively, our data show that PPT mutant cells exhibit loss of oxidative mitochondrial metabolism.

We next asked which mitochondrial target(s) of PPT were most prominently responsible for the loss of mitochondrial oxidative function. As mentioned above, there are three known 4’PP-modified proteins in mammalian mitochondria, NDUFAB1, ALDH1L2, and DHRS2^1^. To explore which of these putative targets displayed the strongest genetic relationships to PPT, we analyzed gene co-dependency relationships from the DepMap Public 23Q4+ Score Chronos dataset^10^ to identify gene functionality within the same biological pathway^11^. Among the three known mitochondrial 4’PP-modified proteins, *AASDHPPT* (PPT) gene effect was positively correlated only with *NDUFAB1* (Pearson r = 0.24) (Extended Data Fig. 2A).

Unique among 4’PP-modified proteins, NDUFAB1 is a structural subunit of complex I of the ETC, as well as an integral part of the mitochondrial fatty acid synthesis (mtFAS) pathway. mtFAS builds *de novo* fatty acyl chains on NDUFAB1, which are required both as precursors for the biosynthesis of lipoic acid (LA), an important TCA cycle enzyme co-factor, as well as to stabilize ETC assembly factors and iron-sulfur (FeS) cluster biogenesis complexes via protein-protein interactions with NDUFAB1^12,9^. Because the newly synthesized fatty acids are added to the terminal sulfhydryl group of the 4’PP PTM, the addition of a 4’PP to NDUFAB1 is required for mtFAS to occur and to support these downstream pathways^13^. Of note, the mtFAS pathway gene *OXSM* ranked third overall for co-dependency with *AASDHPPT* (Pearson r = 0.42)^14^ out of all genes in the Depmap database (Extended Data Fig. 2B). Additional mtFAS pathway genes also ranked highly in terms of *AASDHPPT* co-dependency, including *MCAT* and *MECR* (Pearson r = 0.36 and 0.32 respectively) (Extended Data Fig. 2C and D). Given that the 4’PP modification is necessary for acyl synthesis on NDUFAB1, these data suggest that impaired mitochondrial respiration, reduced levels of ETC complexes, and altered TCA cycle metabolism in PPT-deficient cells may be a downstream result of diminished mtFAS function, in agreement with the known roles of mtFAS^9,15^.

To further explore the genetic relationships between *AASDHPPT* and mtFAS, we performed clustering and two-dimensional mapping of pairwise gene essentiality score correlations with *AASDHPPT*. We included genes associated with the GO term “pantothenate metabolism” from the DepMap project database as well as all genes encoding known phosphopantetheinylated proteins, mtFAS pathway genes, and genes involved in downstream lipoic acid synthesis and ETC assembly -such as the leucine, t<u>y</u>rosine, arginine <u>m</u>otif (LYRM) assembly factors which are regulated by mtFAS^9^. The analyzed genes grouped into four distinct clusters based on gene essentiality (Extended Data Fig. 2E). Two-dimensional mapping of correlation distance showed that *AASDHPPT* most strongly correlated with mtFAS pathway genes, as well as several LYRM family members, further suggesting that PPT may act as a critical regulator of mtFAS functional activity (Fig. 2A).

**Figure 2.**
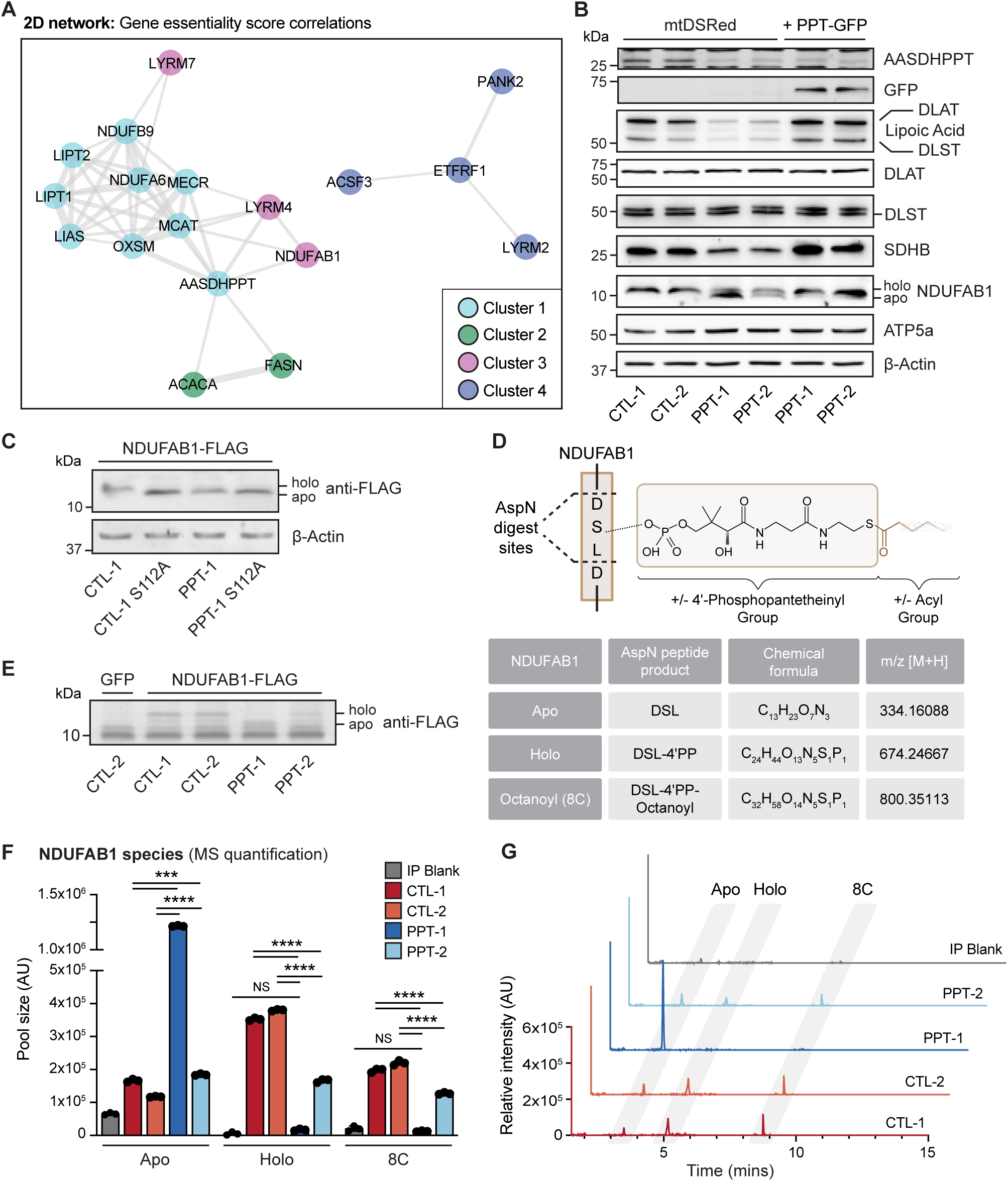
AASDHPPT is required for mitochondrial fatty acid synthesis pathway function. (A) Two-dimensional network diagram representing gene essentiality score correlations between selected genes associated with *AASDHPPT* and known substrate targets, mtFAS pathway members, LYRM family members, and the GO term “pantothenate metabolism” using a minimum threshold correlation score > 0.2. Different colors represent genes associated with clusters as shown in Extended Data Fig. 2E. Correlation strength is represented by the length and thickness of the connecting edge. (B) Whole cell lysates from indicated cell lines expressing mtDSRed or PPT-GFP fusion protein were separated by SDS-PAGE and immunoblotted for the denoted targets. Data are representative of 3 biological replicates. Red asterisk denotes ATP5a band from previous blot. (C) Whole cell lysates from indicated cell lines expressing either NDUFAB1-Flag or NDUFAB1_S112A_-Flag were separated by SDS-PAGE and immunoblotted for protein expression of the denoted targets. Data are representative of 3 biological replicates. (D) Schematic of mass spectrometry approach using AspN digestion to detect Apo-, Holo-, and Acyl-species of NDUFAB1. (E) Coomassie stain of SDS-PAGE gel of NDUFAB1-FLAG immunoprecipitation from the indicated cell line. (F) Relative quantification of NDUFAB1 AspN digest products via tandem liquid chromatography mass spectrometry n=3 technical replicates from immunoprecipitation, representative of 3 distinct biological replicates. NS = not significant, *** = p<0.001, **** = p<0.0001 as determined by two-way ANOVA followed by Šidák’s multiple comparisons test. (G) Representative overlayed extracted ion chromatograms of Apo-, Holo-, and Acyl-NDUFAB1 from cells of the indicated genotype, quantified in F.

To test this hypothesis, we examined whether PPT-deficiency altered known mtFAS pathway endpoints, including protein lipoylation and succinate dehydrogenase subunit B (SDHB) stability, which is a reliable surrogate of the LYRM-dependent ETC assembly mechanism we previously described^9^. PPT-deficient cells exhibited reduced lipoylation of pyruvate dehydrogenase (PDH) and ⍺-ketoglutarate dehydrogenase (OGDH) subunits (DLAT and DLST, respectively) along with reduced stability of SDHB (Fig. 2B), resembling the molecular phenotype of mtFAS-deficient cells. These findings also agree well with our metabolic tracing data, which suggest reduced TCA cycle activity downstream of SDH (Fig. 1H, Extended Data Fig.1F-H). PDH and OGDH lipoylation and SDHB stability were all rescued by transient re-expression of PPT-GFP compared to a control vector expressing mitochondrially localized DSRed (mtDSRed) (Fig. 2B).

Although one prior study demonstrated that PPT is able to add a 4’PP group to NDUFAB1 *in vitro*^2^, its requirement for NDUFAB1 phosphopantetheinylation had never been tested in mammalian cells. We noted that NDUFAB1 migrated faster on SDS-PAGE gels in PPT-deficient cell lysates (Fig. 2B, Extended Data Fig. 2G), and hypothesized that this difference in migration might correspond to loss of the 4’PP-modification, as unmodified apo-NDUFAB1 has previously been shown to migrate faster than 4’PP-modified holo-NDUFAB1 due to its lower molecular weight^2^. We asked whether this downward shift in NDUFAB1 migration corresponded to the 4’PP modification by mutating the modified serine to an alanine residue (S112A), thereby blocking phosphopantetheinylation of NDUFAB1. We expressed wild type NDUFAB1-Flag or NDUFAB1_S112A_-Flag in both control and PPT-deficient cells and found that NDUFAB1_S112A_-Flag in control cells migrates at the same molecular weight as NDUFAB1-Flag in PPT-deficient cells (Fig. 2C). Moreover, NDUFAB1_S112A_-Flag does not display any further downward shift in PPT-deficient cells, indicating that the two perturbations are not additive, and are both likely to be due to loss of 4’PP at S112 (Fig. 2C).

To directly assess whether phosphopantetheinylation of NDUFAB1 is decreased in PPT-deficient cells, we developed a method to detect 4’PP-modification of NDUFAB1 by liquid chromatography mass spectrometry (LCMS), borrowing from an assay developed in plant mitochondria and recently demonstrated for the first time in mammalian cell lines (Fig. 2D)^16,17^. We expressed and immunoprecipitated NDUFAB1-Flag in control and PPT-deficient cells and found that our isolated NDUFAB1 exhibited a similar gel shift as we had observed by blotting in whole cell lysates (Fig. 2E). We then digested these samples with an aspartate endoproteinase (AspN), resulting in a tripeptide (Asp-Ser-Leu) that we could monitor for 4’PP- or 4’PP-acyl modifications via LCMS. This allowed us to detect and quantify levels of apo-, holo-, and octanoyl-NDUFAB1 from control and PPT-deficient cells, verified by standards synthesized *in vitro* (Extended Data Fig. 2H). We found that relative to control cells, PPT-deficient cells have reduced levels of both holo- and octanoyl-NDUFAB1 (Fig. 2F and G). The PPT-1 clone displayed a striking accumulation of apo-NDUFAB1, in agreement with its stronger loss of function phenotype. In addition to the loss of holo-NDUFAB1, we found a modest increase in the levels of intracellular CoA in the PPT-1 and PPT-2 cell lines (Extended Data Fig. 2F). From these data, we conclude that PPT is required for 4’PP modification of NDUFAB1 and downstream mtFAS activity, including protein lipoylation, ETC assembly, and mitochondrial respiration, which all require mtFAS products^15^.

The question then became: if PPT is required for modification of mitochondrial proteins, does it act in the cytoplasm, prior to mitochondrial import, or is there an undescribed mitochondrial pool of PPT? To address this question, we performed immunofluorescence staining for endogenous PPT and mitochondrial ATP synthase subunit 5a (ATP5a) and assessed co-localization (Fig. 3A and B, Extended Data Fig. 3A-C). As expected, PPT expression was significantly reduced in PPT-deficient cells compared to controls (Extended Data Fig. 3B) and exhibited a punctate, perinuclear expression pattern that co-localized with ATP5a staining in control cells (Pearson r = 0.60 and 0.62) (Fig. 3B). Additionally, signal in the PPT channel was surprisingly observed in the nuclei of some cells (Fig. 3A, Extended Data Fig. 3A).

**Figure 3.**
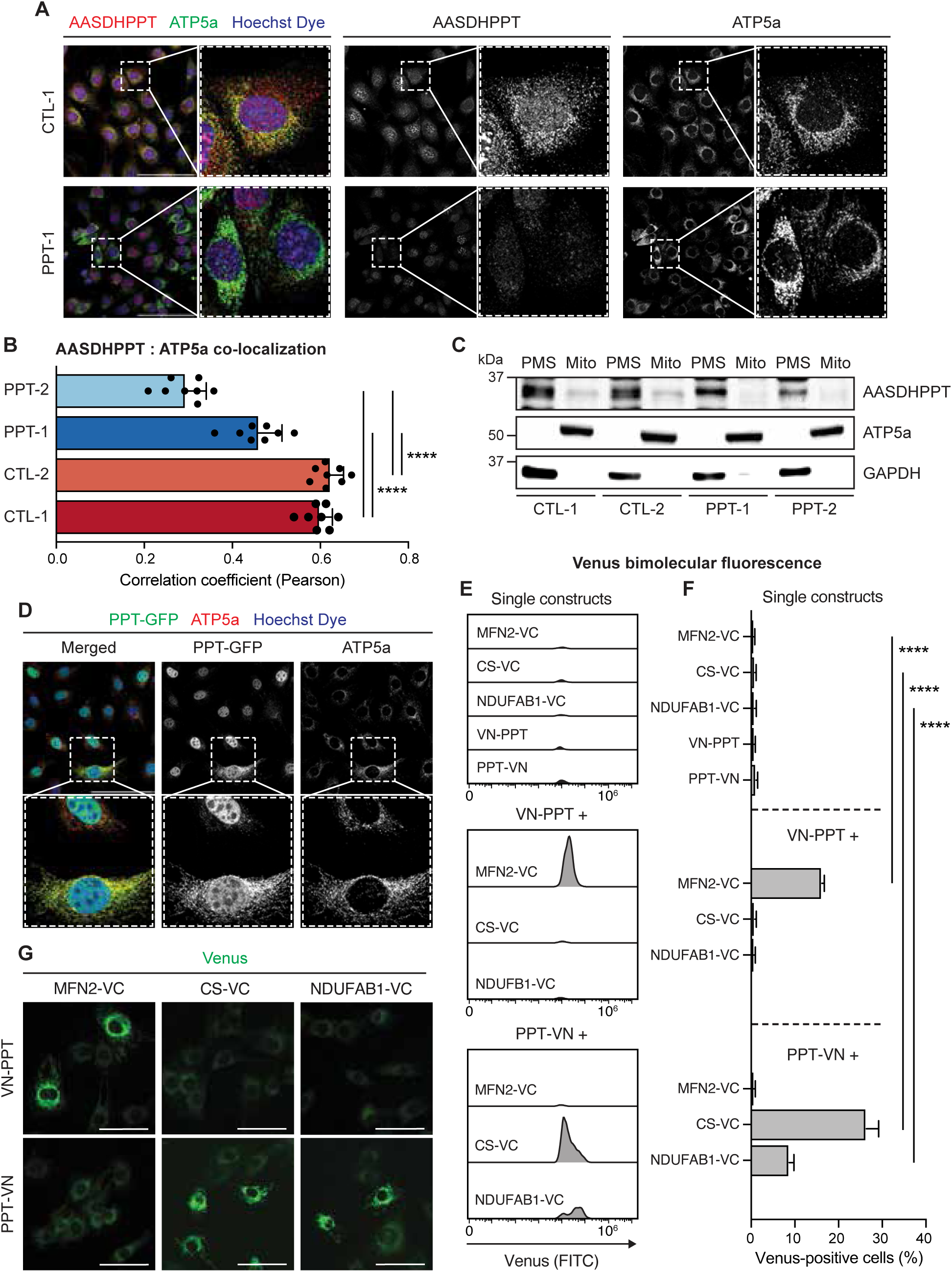
AASDHPPT localizes to the mitochondrial matrix. (A) Representative 40X HPF images of the indicated cell lines immunostained with Hoechst dye to visualize nuclei and antibodies against AASDHPPT (red) and ATP5a (green). Scale bars = 100µm. (B) Quantification of AASDHPPT and ATP5a signal colocalization outside of the nucleus from n = 8 40X HPF per group assessed by Pearson’s correlation coefficient. (C) Subcellular fractionation of *Aasdhppt* mutant cell lines and control. Post-mitochondrial supernatant (PMS) and mitochondrial lysate (Mito) was isolated from the indicated cell lines and immunoblotted for denoted targets. Data are representative of 3 biological replicates. (D) Representative 40X HPF images of WT skeletal muscle myoblasts expressing AASDHPPT-GFP (PPT-GFP) stained with Hoechst dye and antibody against ATP5a (red). Scale bars represent 100 μm. Dashed outlines denote magnified regions shown in panels with solid outlines. For all statistical analysis data represent the mean; error bars denote SD. **** = p<0.0001 as determined by one-way ANOVA with Šidák’s multiple comparisons test. (E) Representative histograms of Venus (FITC) fluorescence in the indicated BiFC cell lines. (F) Percentage of Venus-positive cells in the indicated cell lines as measured by flow cytometry. Data represent the mean; error bars denote standard deviation (n = 3 per group). Statistical significance was determined by one-way ANOVA with Tukey’s multiple comparison test. (G) Representative 20X images of endogenous Venus fluorescence in VC + VN BiFC cell lines. Scale bars represent 50 μm.

To test whether this staining pattern was specific, we employed siRNA and found a similar reduction in endogenous PPT signal as observed in our PPT-deficient cell lines, although nuclear signal in the PPT channel was maintained despite knockdown (Extended Data Fig. 3D). We therefore also expressed PPT-Flag and performed immunofluorescent staining for the Flag epitope and found a similar punctate, peri-nuclear staining pattern that overlapped with citrate synthase staining but displayed no clear nuclear signal (Extended Data Fig. 3E). These results indicate that a sub-population of PPT localizes specifically to mitochondria but not to the nucleus.

We further verified the observed mitochondrial localization using subcellular fractionation and Western blotting in control and PPT-deficient cells. We detected endogenous PPT expression in both mitochondria and post-mitochondrial supernatant (PMS) (Fig. 3C). PPT expression was reduced in both fractions, and nearly absent from mitochondria in PPT-deficient cells (Fig. 3C). We also characterized subcellular localization of PPT-GFP using immunofluorescence and found strong signal overlap with ATP5a (Fig. 3D), in agreement with our prior observations.

We then asked whether this pool of mitochondrially-localized PPT was mitochondrially associated or in the mitochondrial matrix with NDUFAB1 and the rest of the mtFAS machinery. To tackle this question, we employed a bimolecular fluorescence complementation (BiFC) system^18^ to visualize the sub-mitochondrial localization of PPT. We used a split-Venus tag technique in which the N-terminus of the Venus fluorophore (VN, residues 1-172) was attached to either the N- or C-terminus of PPT, while the C-terminus of Venus (VC, residues 155-238) was attached to the C-terminus of a control protein with known localization. This system allows us to observe fluorescent signal only when both halves of the split-Venus tag are able to associate within a compartment^18^. For these control proteins, we chose Mitofusin 2 (MFN2) – a transmembrane protein that localizes on the outer mitochondrial membrane with both its N-and C-termini facing the cytoplasm – along with citrate synthase (CS) and NDUFAB1, both of which are matrix-localized. We quantified Venus fluorescence in each of our BiFC conditions by flow cytometry, and observed significant Venus signal only when PPT-VN cells were co-expressed with either CS-VC or NDUFAB1-VC, indicating that PPT localizes to the mitochondrial matrix (Fig. 3E-G, see Extended data Fig. 3F for gating strategy).

Most mitochondrial proteins are encoded by nuclear genes and imported into mitochondria via an N-terminal mitochondrial targeting sequence (MTS), consisting of approximately 20 to 40 amino acid residues^19^. Surprisingly, in cells expressing N-terminally tagged PPT (VN-PPT), positive Venus fluorescence was only detected when co-expressed with MFN2-VC (Fig. 3E-G), indicating that N-terminally tagged VN-PPT is mitochondrially associated on the OMM. Since our BiFC data showed that tagging PPT on the N-terminus prevented its localization in the mitochondrial matrix (Fig. 3E-G), we hypothesized that mitochondrial localization of PPT may be driven by an N-terminal MTS.

To explore this hypothesis, we first analyzed the AASDHPPT coding sequence using the MitoFates^20^ and DeepMito^21,22^ prediction algorithms. MitoFates reports a 41.5% probability of a mitochondrial presequence at the N-terminus of AASDHPPT, with a TOM20 recognition motif from amino acids 5-9 and an MPP cleavage site at amino acid 19 (W) (Fig. 4A and B). Alternatively, DeepMito predicts a 47% probability of mitochondrial localization. This region of the amino acid sequence is well conserved among mammals (Extended Data Fig. 4A). To test whether the N-terminus of PPT is a bonafide MTS, we generated N-terminal truncations to the next two methionine residues at positions 14 and 37 (PPT_∆1-13_-GFP and PPT_∆1-36_-GFP, respectively, Fig. 4A) and examined their localization using fluorescence microscopy. While PPT-GFP co-localized strongly with ATP5a, PPT_∆1-13_-GFP exhibited diffuse localization throughout the nucleus and cytoplasm (Fig. 4C-D). PPT_∆1-36_-GFP exhibited weak fluorescent expression mostly within the nucleus, suggesting that this larger N-terminal truncation destabilizes PPT (Fig. 4C-D). To further refine the MTS based on the MitoFates prediction, we also tested an N-terminal truncation replacing the first 20 amino acids with a single methionine (PPT_∆1-20_-GFP). PPT_∆1-20_-GFP behaved similarly to PPT_∆1-36_-GFP, with very low levels of diffuse expression (Extended Data Fig. 4B), perhaps indicative of instability upon completely failed import. Sub-cellular fractionation and Western blotting supported our microscopy results, showing that while PPT-GFP and PPT_∆1-13_-GFP are both present in PMS and mitochondrial fractions, PPT_∆1-13_-GFP accumulates in the PMS (Fig. 4E), suggesting difficulty in localizing to mitochondria.

**Figure 4.**
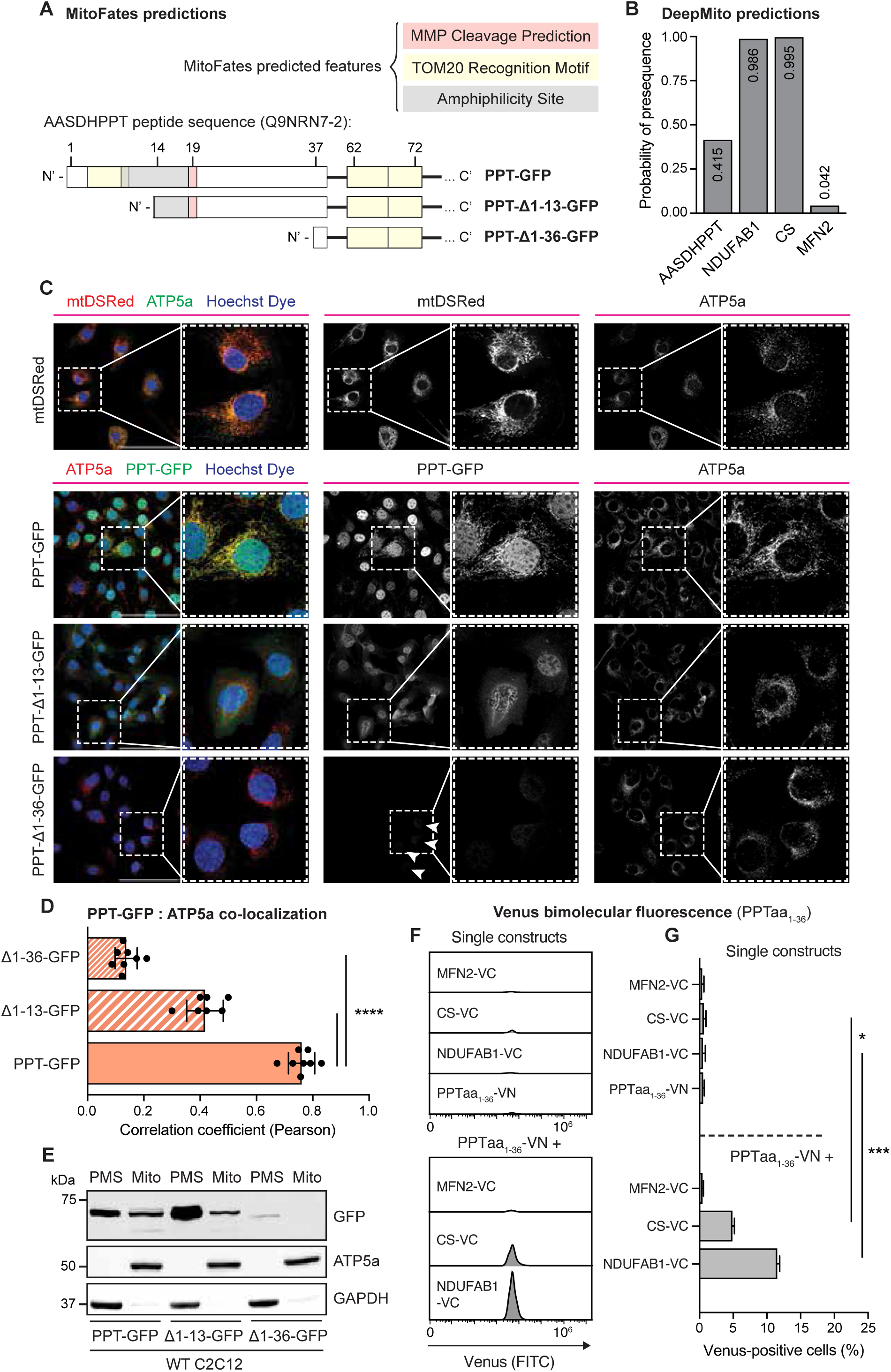
AASDHPPT has a N-terminal MTS. (A) Schematic Illustration of MitoFates^20^ predicted features in full length AASDHPPT (PPT-GFP) and AASDHPPT with a 1-13 (∆1-13) or 1-36 (∆1-36) N-terminal truncation. (B) Quantification of MTS presequence probability in AASDHPPT, NDUFAB1, Citrate Synthase (CS) and Mitofusin 2 (MFN2) determined by MitoFates. (C) Representative 40X HPF images of WT cells expressing mtDSRed, PPT-GFP, 13 residue N-terminal truncation of PPT (PPT_∆1-13_-GFP), or 36 residue N-terminal truncation of PPT (PPT_∆1-36_-GFP) constructs, stained with Hoechst dye and antibody against ATP5a (red). Scale bars represent 100 μm. Arrows denote weak expression of GFP signal detected. (D) Quantification of PPT-GFP and ATP5a signal colocalization outside of the nucleus from n = 8 40X HPF assessed by Pearson’s correlation coefficient from experiment shown in (C). Data represent the mean; error bars denote SD. **** = p<0.0001, determined by one-way ANOVA with Dunnett’s multiple comparisons test. (E) Subcellular fractionation of WT cells expressing PPT-GFP, PPT_∆1-13_-GFP, or PPT_∆1-36_-GFP. Post-mitochondrial supernatant (PMS) and mitochondrial lysate (Mito) was isolated from the denoted cell lines, separated by SDS-PAGE, and immunoblotted for the indicated targets. Data are representative of 3 biological replicates. (F) Representative histograms of Venus (FITC) fluorescence in the indicated BiFC cell lines. (G) Percentage of Venus-positive cells in the indicated cell lines as measured by flow cytometry. Data represent the mean; error bars denote standard deviation (n = 3 per group). Statistical significance was determined by one-way ANOVA with Tukey’s multiple comparison test. **** = p<0.0001

We also performed the converse experiment, in which we added only the first 36 amino acids of PPT to GFP, to test the sufficiency of the putative MTS to drive GFP to the mitochondria (PPT MTS-GFP). PPT MTS-GFP showed strong overlap with ATP5a staining by fluorescence microscopy as well as strong expression in the mitochondrial fraction via Western blot (Extended Data Fig. 4C and D). We took a parallel BiFC approach using the split-Venus tag system by expressing the first 36 amino acids of PPT fused to the N-terminus of Venus (PPTaa_1-36_-VN) with either MFN2-VC, CS-VC, or NDUFAB1-VC. As with C-terminally tagged PPT-VN, significant Venus signal was only observed when PPTaa_1-36_-VN was co-expressed with either CS-VC or NDUFAB1-VC (Fig. 4F and G, see Extended Data Fig. 3F for gating strategy), indicating that the first 36 amino acids of PPT are sufficient to target proteins to the mitochondrial matrix.

To test whether this newfound mitochondrial localization is required for PPT to modify mitochondrial target proteins, we transduced PPT-deficient cells with mtDSRed, PPT-GFP, PPT_∆1-13_-GFP, or PPT_∆1-36_-GFP, and assessed mtFAS pathway activity. Expression of PPT-GFP rescued protein lipoylation and SDHB stability, whereas PPT_∆1-13_-GFP and PPT_∆1-36_-GFP both failed to rescue either of these mtFAS-dependent endpoints, nor did they rescue the gel shift of NDUFAB1 (Fig. 5A, Extended Data Fig. 5A). These data demonstrate that mitochondrial localization of PPT is required for 4’PP-modification of NDUFAB1. To further investigate whether mitochondrial localization of PPT is required for ETC assembly and OXPHOS activity, we performed BN-PAGE and Seahorse mitochondrial stress tests as before with control and PPT-deficient cells expressing mtDSRed, PPT-GFP, or PPT_∆1-13_-GFP. We found that ETC complex levels, along with basal and maximal oxygen consumption rates, were rescued by expression of PPT-GFP, and that PPT_∆1-13_-GFP rescued less well than the full length PPT-GFP construct (Fig. 5B-D, Extended Data Fig. 5B-D).

**Figure 5.**
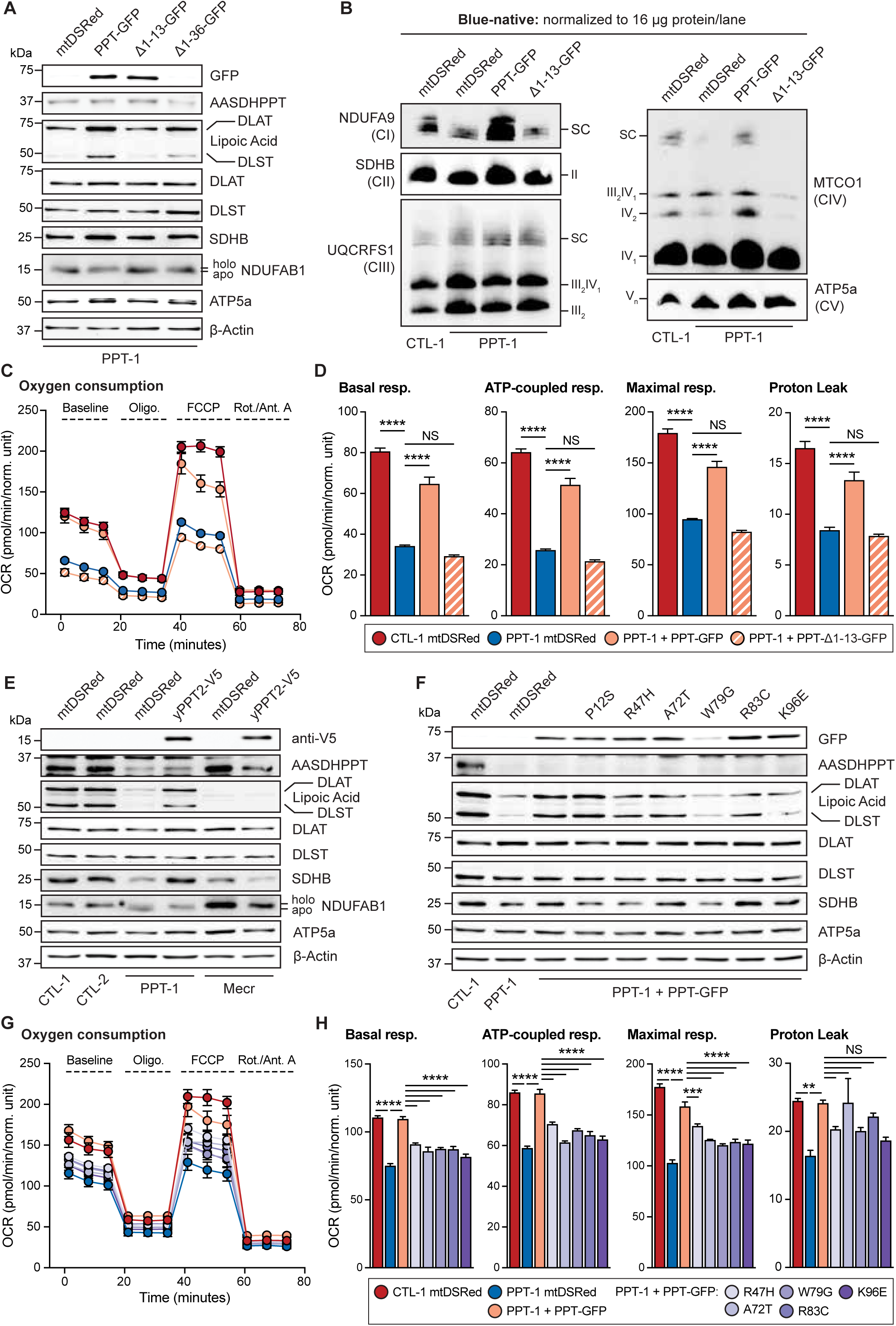
Mitochondrial localization of AASDHPPT is required for mtFAS function. (A) Whole cell lysates from cells expressing mtDSRed, PPT-GFP, PPT_∆1-13_-GFP, or PPT_∆1-36_-GFP were separated by SDS-PAGE and immunoblotted for the denoted targets. Data are representative of n > 3 biological replicates. (B) Blue-native PAGE separation of isolated mitochondrial protein complexes from denoted cell lines expressing mtDSRed, PPT-GFP, or PPT_∆1-13_-GFP, followed by immunoblot with the indicated antibodies. Data are representative of n > 3 biological replicates. (C) Seahorse mitochondrial stress test for Oxygen Consumption Rate (OCR) of indicated cell lines expressing mtDSRed, PPT-GFP, or PPT_∆1-13_-GFP. Data are representative of 1 experiment from 3 biological replicates. Error bars represent SD. (D) Quantification of basal respiration, ATP-production coupled respiration, maximal respiration and proton leak from data in (C). Data are mean OCR; error bars are SEM. **** = p <0.0001, determined by one-way ANOVA with Tukey’s multiple comparisons test. (E) Whole cell lysates from clonal control, *Aasdhppt-*, and *Mecr-*deficient cells expressing mtDSRed or yPpt2-V5 were separated by SDS-PAGE and immunoblotted for the denoted targets. Data are representative of 3 biological replicates. (F) Whole cell lysates from the indicated cell lines expressing mtDSRed, PPT-GFP P12S, PPT-GFP R47H, PPT-GFP A72T, PPT-GFP W79G, PPT-GFP R83C, or PPT-GFP K96E were separated by SDS-PAGE and immunoblotted for the denoted targets. Immunoblots are representative of 3 biological replicates. (G) Seahorse mitochondrial stress test for Oxygen Consumption Rate (OCR) of indicated cell lines expressing mtDSRed, PPT-GFP, PPT-GFP R47H, PPT-GFP A72T, PPT-GFP W79G, PPT-GFP R83C, or PPT-GFP K96E. Data are representative of 1 experiment from 3 biological replicates. Error bars represent SD. (H) Quantification of basal respiration, ATP-production coupled respiration, maximal respiration and proton leak from data in (G). Data are mean OCR; error bars are SEM. ** = p <0.01, *** = p < 0.001, **** = p <0.0001, determined by one-way ANOVA with Šidák’s multiple comparisons test.

Next, we took advantage of the *Saccharomyces cerevisiae* mitochondrial PPTase, yPpt2, (which is known to specifically reside in the matrix) to test if 4’PP mitochondrial modifications rescue the mtFAS-associated mitochondrial defects observed in PPT mutant cells. Indeed, yPpt2-V5 localizes strongly to mitochondria (Extended Data Fig. 5E) and efficiently restores protein lipoylation, SDHB levels, and NDUFAB1 migration in PPT-deficient cells (Fig. 5E). Notably, yPpt2-V5 expression does not rescue protein lipoylation or SDHB in cells deficient for the mtFAS enzyme MECR, indicating that this rescue is specific to loss of PPT and cannot compensate for direct impairments to mtFAS itself (Fig. 5E, Extended data Fig. 5F). Together with our truncation mutant analyses, these data demonstrate that mitochondrial localization of PPT is necessary and sufficient for phosphopantetheinylation of NDUFAB1, mtFAS pathway activity, ETC assembly and mitochondrial respiration.

We recently described a pathogenic variant in the mtFAS gene *MCAT* c.812T>C; p.T271I (rs760294168; NM_173467.4)^23^. As our data demonstrate that perturbed *Aasdhppt* expression results in similar phenotypes as mtFAS loss, we wondered whether there may be patients with pathological variants in *AASDHPPT*. We searched the ClinVar database^24^ and identified several missense variants of uncertain significance (VUS) in *AASDHPPT* that were associated with suspected inborn genetic disease. We selected six of these variants for analysis: c.343C>T; p.P12S (NM_015423.3), c.140G>A; pR47H (NM_015423.3), c.214G>T; pA72T (NM_015423.3), c.235T>G; pW79G (NM_015423.3), c.247C>T; pR83C (NM_015423.3), and c.286A>G; pK96E (NM_015423.3).

We created each of these missense mutations in our full length PPT-GFP construct and assessed their ability to rescue perturbed mtFAS activity in our PPT-deficient cells. We found that all variants, with the exception of PPT-GFP P12S, exhibited weaker rescue effects than wild type PPT (Fig. 5F, Extended Data Fig. 5G). These variants of PPT were also unable to completely rescue basal, ATP-production coupled, or maximal respiration rates compared to wild type PPT-GFP (Fig. 5G and H). Of note, cells expressing the PPT-GFP W79G variant showed low levels of protein expression compared to the other variants (Fig. 5F, Extended Data Fig. 5G-H) despite expression from the same promoter, suggesting that this variant may result in protein instability. These data identify R47H, A72T, W79G, R83C, and K96E as potential clinically observed pathologic variants of *AASDHPPT*, with molecular phenotypes attributable to altered mtFAS activity and perturbed cellular respiration.

Our data position PPT as a critical mediator of mitochondrial oxidative function, due to its requirement for proper function of the mtFAS pathway. While we show definitively that mitochondrial PPT is required for phosphopantetheinylation of NDUFAB1, we did not explore its involvement in 4’PP-modification of ALDH1L2 and DHRS2 in the present study. Future work focused on the role of PPT in these additional pathways will be important to expand our understanding of the full scope of PPT action in cells. The discovery of distinct pools of PPT raises questions about what factors control PPT activity in each compartment. For instance, PPT uses CoA as the substrate for 4’PP-modification of target proteins. How perturbations in CoA availability affect PPT enzymatic activity may have important implications for PPT targets such as mtFAS in diseases with CoA dysregulation^25^. Moreover, our data demonstrate that PPT is targeted to mitochondria via an N-terminal MTS, but how this localization is controlled in cells to produce two distinct pools is completely unstudied. Whether PPT localization is dynamic will also be an interesting and important area for future study. Of note, we identified five probable pathogenic patient variants in *AASDHPPT*. Our results predict that patients harboring these mutant alleles likely exhibit impaired mtFAS, leading to diminished mitochondrial function. These data add to the growing list of patients with pathogenic variants in mtFAS and lipoic acid synthesis genes and support the notion that impairments in CoA metabolism may cause disease via mtFAS dysfunction^26,27^.

## Methods

### Aasdhppt mutant cell lines, siRNA transfection, growth assays, and viral transduction Cell Culture

C2C12 immortalized mouse skeletal myoblasts (ATCC CRL-1772, verification provided by ATCC) were grown in DMEM supplemented with 4.5 g/L glucose, glutamine, and sodium pyruvate (Corning, 10-013-CV) and 10% FBS (Sigma, F0926) in humidified tissue culture incubators (37°C, 5% CO_2_).

### Generation of Aasdhppt mutant cell lines

Four sgRNA sequences (**Table 1)** were designed targeting exon 1 or exon 2 of the mouse *Aasdhppt* gene and subcloned into the pLentiCRISPR v2 vector using previously described methods^28^. Parental C2C12s were transiently transfected with pLentiCRISPRv2_sgAasdhppt or empty vector using a JetOptimus lipid transfection reagent (Polyplus, 76299-630). 48 hours after transfection, cells were sorted on GFP signal intensity and plated into 96-well plates to obtain single cell colonies for expansion and establishment of stable, clonal cell lines. Clones were screened for AASDHPPT protein expression levels and two control clones and two *Aasdhppt* mutant clones were selected for further experiments. While PPT-deficient clones were derived from both guide RNAs, clones with the strongest loss of PPT expression were all derived from sgRNA 1, and we focused further analyses on these clones with the strongest loss of function phenotypes.

**Table 1.**
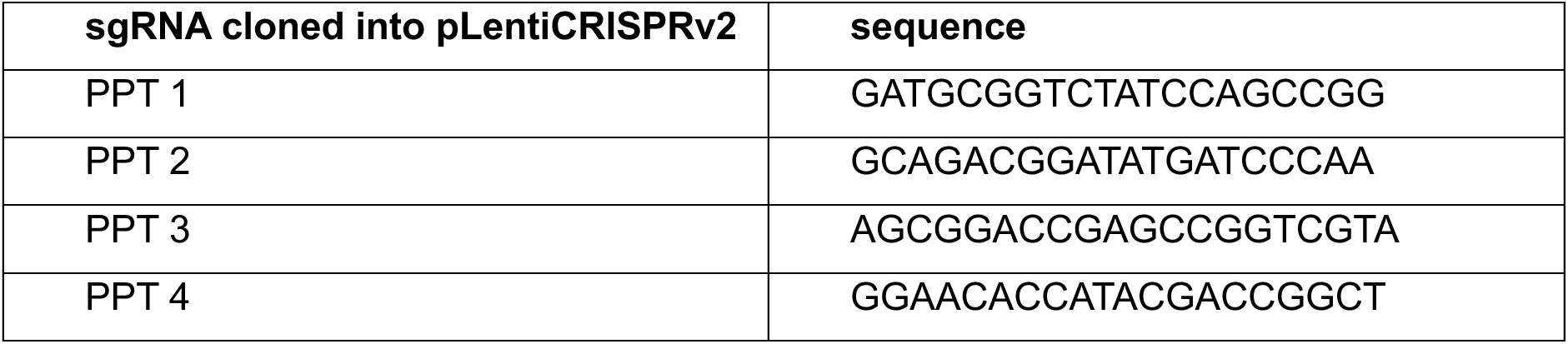

### Production of siRNA knockdown cell lines

To knockdown *Aasdhppt* or *Ndufab1* expression, two Silencer Select pre-designed siRNAs targeting *Aasdhppt* were acquired from Thermo Fisher Scientific and an ON-TARGETplus SMARTpool siRNA targeting *Ndufab1* was acquired from Horizon Discovery (**Table 2**). The siRNAs were packaged with Lipofectamine RNAiMAX transfection reagent for 30 minutes at room temperature according to manufacturer instructions. C2C12 cells were transfected with the siRNA mixture for a final concentration of 40 nM overnight. The following day, cells were provided with fresh growth media and cells were cultured for 72 hours post-transfection to either collect whole-cell lysates for western blot analysis or be fixed in 4% paraformaldehyde for immunofluorescence analysis. C2C12 cells were transfected with ON-TARGETplus Non-targeting Control Pool (D-001810-10-05) as a control.

**Table 2.**
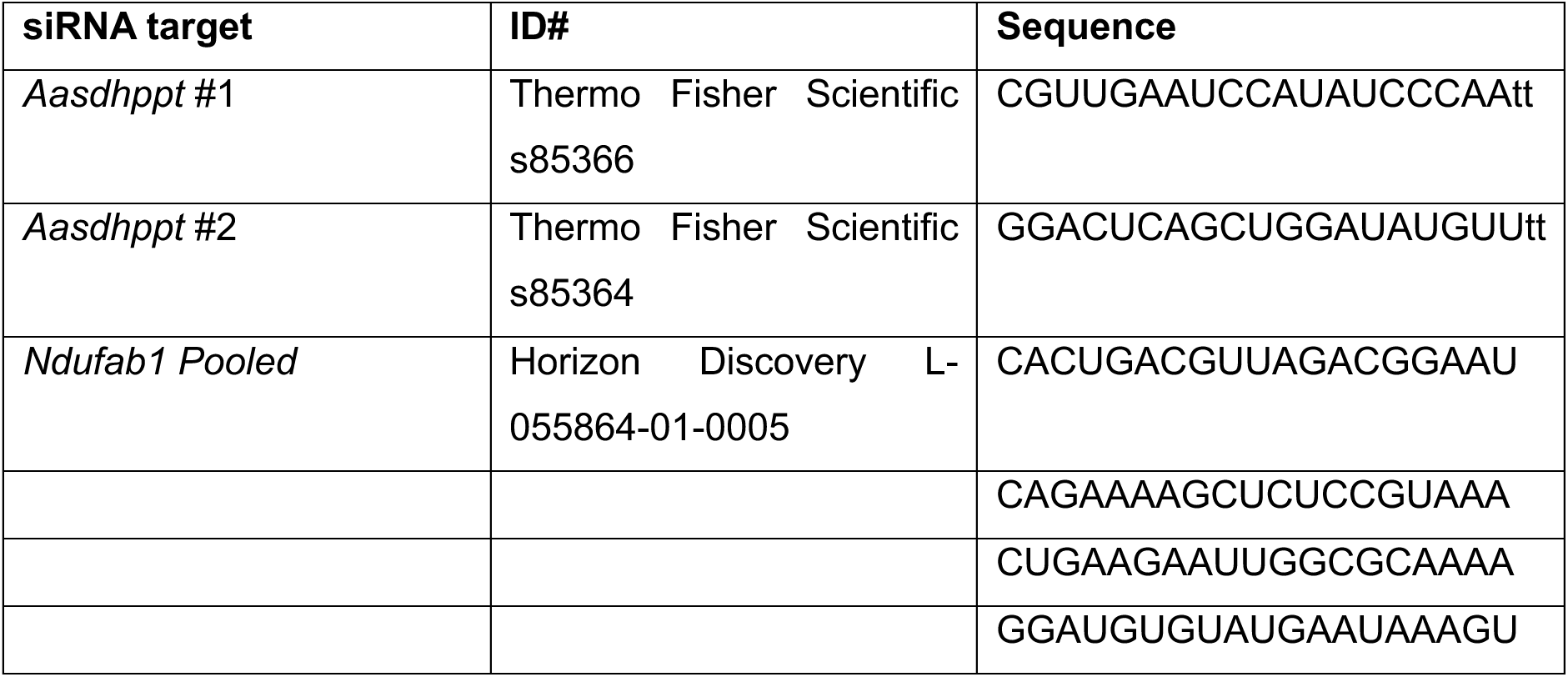

### Retroviral backbone design

For co-localization and rescue experiments, control and *Aasdhppt* mutant clonal lines were transiently infected with retrovirus harboring a control transfer plasmid expressing mtDSRed (pQXCIP mtDSRed) or pMMLV retrovirus vectors custom designed, synthesized, and purchased from VectorBuilder (Chicago, IL, USA), harboring full length, or N-terminal truncations of residues 1-13 or 1-36, of AASDHPPT with C-terminal GFP and Flag tag-fusion, or yPPT2 with C-terminal Flag tag-fusion. *<u>Site-directed mutagenesis.</u>* For generation of retrovirus constructs harboring single point mutations in the full length AASDHPPT-GFP-Flag construct, primers in **Table 3** were used with a New England Biolabs Q5 Site-Directed Mutagenesis Kit. Plasmid reactions were transformed into competent *E. coli* and selected colonies were sequence validated for successful incorporation of point mutations.

**Table 3.**
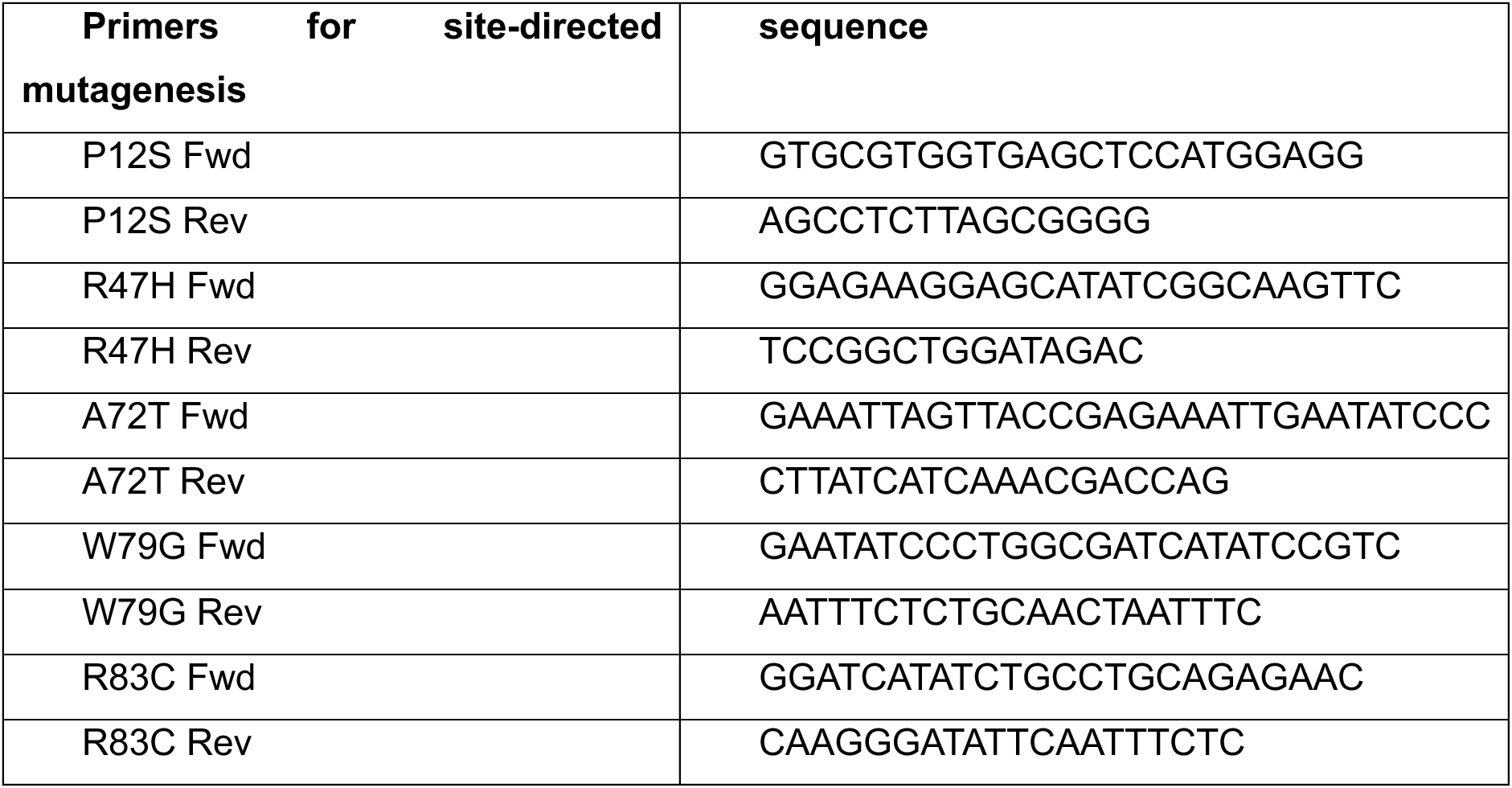
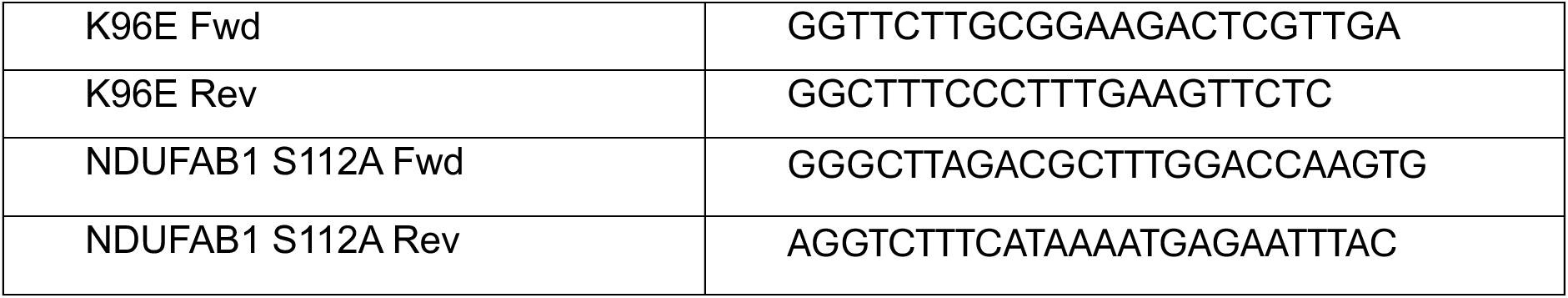

### Growth Assays

For growth assays 5,000 cells/well were plated in 24-well plates and grown in DMEM with either 10% FBS, 4 mM glutamine, and 25 mM glucose for 4 days or 20% FBS, 4 mM glutamine, and 10 mM galactose for 2 days in technical quadruplicates. Tissue culture plates were placed in an IncuCyte® chamber (Sartorius) and percent confluency was measured from sixteen images at 10X magnification per well every 3 hours. Percent confluency curves from technical replicates were fit with a non-linear regression during the logarithmic growth phase using the exponential growth equation in GraphPad Prism to determine population doubling time.

### Genomic Sequence Analysis of AASDHPPT Mutants

Approximately five million cells of each clonal mutant cell line, PPT-1 and PPT-2, were pelleted and washed with PBS then frozen at -80°C. Genomic DNA was extracted using a column-based kit assay (Qiagen 69504). Aasdhppt was amplified using primers described in **Table 4**, and sanger sequenced using the forward primer. Sanger sequences were analyzed using Snapgene software.

**Table 4.**
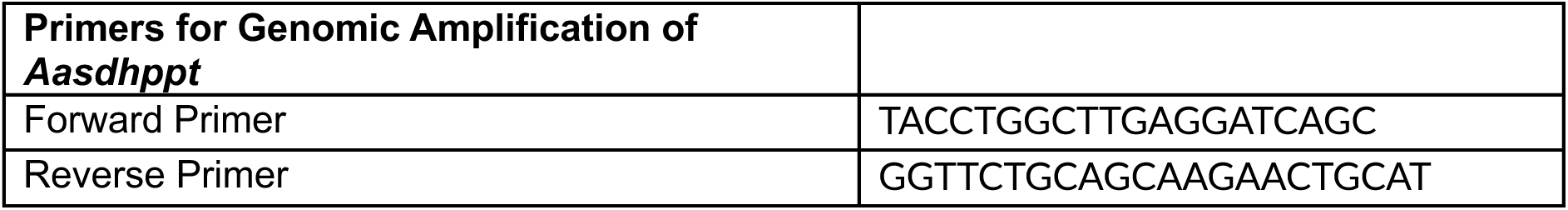

### Crude mitochondrial isolation

Cells were harvested and processed as previously described^9^. Briefly, cells were harvested by trypsinization, washed once with cold, sterile PBS (Gibco, 10010-023), then stored at -80°C. At the time of mitochondrial isolation, cell pellets were thawed and resuspended in 1 mL CP-1 buffer (100 mM KCl, 50 mM Tris-HCL, 2 mM EGTA, pH 7.4) supplemented with mammalian protease inhibitor cocktail (mPIC, Millipore Sigma P8340) and mechanically lysed by 10 passes through a 1 mL insulin syringe with 26-guage needle and centrifuged at 700 x g for 10 minutes to pellet unlysed cells and debris. Supernatant was transferred to a new microcentrifuge tube and centrifuged at 10,000 x g for 10 minutes to pellet the crude mitochondrial fraction. Post-mitochondrial supernatant (PMS) was used for Western blot analysis or discarded. Mitochondrial pellets were resuspended in a small volume of either RIPA buffer (10 mM Tris-HCl, pH 8.0, 1 mM EDTA, 0.5 mM EGTA, 1% Triton X-100, 0.1% Sodium Deoxycholate, 0.1% SDS, 140 mM NaCl) supplemented with mPIC, or CP-1 buffer, equal to approximately twice the pellet volume. Resuspended pellets were then used for applications described below.

### SDS-PAGE and immunoblotting

Cultured cells were scraped from tissue culture plates directly into RIPA buffer supplemented with mammalian protease inhibitor cocktail (mPIC, Millipore Sigma) and Pierce Universal Nuclease, incubated at 4°C with constant agitation on a microcentrifuge tube orbital shaker, then centrifuged at 17,000 x g for 20 minutes at 4°C to remove insoluble material. Supernatant was then transferred to a new microcentrifuge tube to be saved as whole cell lysate (WCL). For protein analysis, WCL, PMS, and/or crude mitochondrial fractions were first normalized for total protein content via Pierce BCA Protein assay (Thermo Fisher Scientifc 23225). Samples were resolved by SDS-PAGE and transferred to nitrocellulose membranes. Immunoblotting was performed using the indicated primary antibodies (**Table 5**) with visualization and analysis of protein signal using Rabbit and Mouse IgG Antibody DyLight 680, or 800, Conjugated secondary antibodies and Bio-Rad ChemiDoc Imaging System. Alternatively, secondary anti-mouse or anti-rabbit HRP antibody and SuperSignal West Femto Maximum Sensitivity Substrate (Thermo Fisher Scientific 34096) were used to visualize band signals.

**Table 5.**

| **Antibodies** | **Source and reference number** |
| --- | --- |
| AASDHPPT | Proteintech 11244-1-AP |
| β-actin | Proteintech 66009-1-Ig |
| NDUFA9 | Abcam ab14713 |
| UQCRFS1 | Proteintech 18443-1-AP |
| MTCO1 | Abcam ab110270 |
| ATP5a | Abcam ab14748 |
| Lipoic acid | Millipore 437695 |
| Pyruvate Dehydrogenase E2 (DLAT) | Abcam ab172617 |
| DLST | Cell Signaling 5556 |
| SDHB | Abcam ab14714 |
| NDUFAB1 | Abcam ab96230 and ab181021 |
| GFP | Abcam ab290 |
| Citrate Synthase | Cell Signaling 14309 |
| Anti-V5 | Abcam ab27671 |
| Anti-FLAG | Thermo Fisher MA1-91878 |
| MECR | Proteintech 51027-2-AP |
| Goat anti-Mouse IgG (H+L) AlexaFluor 568 | Thermo Fisher A-11004 |
| Goat anti-Mouse IgG (H+L) Alexafluor 488 | Thermo Fisher A-11029 |
| Goat anti-Rabbit IgG (H+L) AlexaFluor 568 | Thermo Fisher A-11011 |
| Goat anti-Mouse IgG (H+L), HRP | Thermo Fisher A28177 |
| Goat anti-Rabbit IgG (H+L), HRP | Thermo Fisher A16110 |
| Goat anti-Mouse IgG (H+L) Dylight 800 | Rockland Immunochemicals 610-145-002-0.5 |
| Goat anti-Rabbit IgG (H+L) Dylight 800 | Rockland Immunochemicals 611-145-002-0.5 |
| Goat anti-Mouse IgG (H+L) AlexaFluor 680 | Thermo Fisher A-21057 |
| Goat anti-Rabbit IgG (H+L) AlexaFluor 680 | Thermo Fisher A10043 |

### Seahorse Extracellular Flux Rate Assay

Extracellular flux rate assays were performed on a Seahorse XFe96 Analyzer (Agilent). Cells from the indicated clones and experimental conditions were plated in approximately 22 wells of a 96-well seahorse plate at a density of 8,000 (clonal controls) or 10,000 (PPT-deficient clones) cells/well in DMEM +10% FBS and allowed to adhere overnight. The morning of the assay, cells were washed twice and media was replaced with non-buffered seahorse media (Agilent 103680) supplemented with 25mM glucose, 2mM glutamine, and 1mM sodium pyruvate. A standard mitochondrial stress test was then performed using 1μM oligomycin, 3μM FCCP, and 0.5μM Rotenone + 0.5μM Antimycin A. Baseline oxygen consumption was measured three times, followed by acute injection of oligomycin, followed by FCCP and finally Rotenone + Antimycin A. Measurements were standard, with acute injection of drug followed by 3 minutes of mixing then measurements of oxygen levels over 3 minutes, with three measurements per phase. Data were normalized with crystal violet absorption for approximation of cell number or by measurement of well % confluency using an IncuCyte® chamber and a 10X magnification image field per well. Results were analyzed in WAVE software and the Agilent Seahorse Analytics browser-based application for quantification of basal, maximal, ATP-production coupled, and proton leak respiration.

### Blue Native-PAGE

Crude mitochondrial fractions were prepared and normalized for total protein content as previously described. 100 μg of mitochondria were pelleted at 12,500 x g for 5 minutes at 4°C. Supernatant was removed and pellets were resuspended in 54 μL of 1x pink lysis buffer (Invitrogen BN2003). Digitonin (GoldBio D-180-2.5) was added to samples at a final concentration of 1% mass/volume and samples were incubated on ice for 15 min. Insoluble material was pelleted by centrifugation at 17,000 x g for 20 minutes and supernatant was transferred to new microcentrifuge tubes. 3 μL of NativePAGE sample buffer (Invitrogen BN2004) was added to samples and 10 μL of sample (∼16 μg of crude mitochondrial fraction) was loaded per lane on pre-cast 3-12%, Bis-Tris NativePAGE gels (Invitrogen BN1001BOX) with NativePAGE anode buffer (Invitrogen 2001) and dark blue cathode buffer (Invitrogen BN2002) at 150V for 30-40 minutes then switched to light blue cathode buffer and run at 20 V overnight. Gels were then transferred to PVDF membranes (pre-activated in 100% methanol) at 100 V for 2 hours. Following transfer, membranes were washed in 8% acetic acid solution for 5 minutes, washed in sterile distilled water twice for 5 minutes, then washed in 100% methanol thrice for 5 minutes, and finally washed in sterile distilled water thrice for 5 minutes. Membranes were then blocked in 5% BSA solution at room temperature. Following blocking, membranes were incubated with the indicated primary antibodies overnight at 4°C (**Table 5)**. Secondary anti-mouse or anti-rabbit HRP antibody and SuperSignal West Femto Maximum Sensitivity Substrate (Thermo Fisher Scientific 34096) was used to visualize bands with a Bio-Rad ChemiDoc Imaging System.

### Glutamine Tracing

The indicated clonal cell lines were plated in 6-well plates in triplicate at a density of 200K for PPT-1 and PPT-2, and 175K for Ctrl-1 and Ctrl-2 to account for differences in growth rate. Cells were allowed to adhere for two hours, at which point the media was changed to unlabeled media (Corning DMEM 17-207-CV with 1mM sodium pyruvate, 5mM glucose, and 4mM glutamine). After twenty hours in unlabeled media, wells were changed to labeled media at the indicated time points prior to harvest (Corning DMEM 17-207-CV with 1mM sodium pyruvate, 5mM glucose, and 4mM U^13^C-glutamine). Three wells per cell line were left unlabeled for natural isotope abundance correction, and another three per cell line were used to determine cell number at the time of freezing. 24 hours after cells were in unlabeled media, all wells were washed with normal saline once and frozen at -80°C. Plates were then extracted for metabolites using a volume of 40:20:20 methanol:acetonitrile:water scaled to the number of cells determined by the cell count wells (average from three wells). Lysates were sonicated and incubated on ice for 60 minutes. At this point any insoluble material was spun out of the sample at 17,000 x g for 10 minutes at 4°C. 1000 µL of supernatant was dried using a SpeedVac concentrator.

### Liquid chromatography – Mass Spectrometry

Dried metabolomics extracts were next resuspended in 100% water containing 25 µg/mL D5-Glutamate (DLM-556, Cambridge) such that resuspension sample equivalents were

5.82E+3 cells/µL. 2 µL of resuspended samples were injected on column, a process blank was analyzed before and after experimental samples, and a pooled sample was injected twice before experimental samples for column conditioning, and every 7-10 sample injections thereafter. Data dependent MS2 (ddMS2) data was collected from pooled experimental replicates to identify group-specific compounds.

Data were collected on a Vanquish liquid chromatography system coupled to an Orbitrap Exploris 240 (Thermo Fisher Scientific) using a heated electrospray ionization (H-ESI) source in ESI negative mode. All samples were run through a 24-minute reversed-phase chromatography ZORBAX extend-C18 column (1.8μm, 2.1mm × 150mm, 759700-902, Agilent, California, USA) combined with a guard column (1.8μm, 2.1mm × 5mm, 821725-907, Agilent). Mobile phase A was LC/MS H2O (W6-4, Fisher) with 3% LC/MS grade methanol (A456-4, Fisher), mobile phase B was LC/MS grade methanol (A456-4, Fisher), and both mobile phases contained 10 mM tributylamine (90780-100ML, Sigma), 15 mM acetic acid, and 0.01% medronic acid (v/v, 5191-4506, Agilent). For the re-equilibration gradient, mobile phase A was kept the same, and mobile phase B was 99% LC/MS grade acetonitrile (A955-4, Fisher). Column temperature was kept at 35°C, flow rate 0.25 mL/min, and the solvent gradient was as follows: 0-2.5 min held at 0% B, 2.5-7.5 min from 0% B to 20% B, 7.5-13 min from 20% B to 45% B, 13-20 min from 45% B to 99% B, and 20-24 min held at 99% B. The analytical solvent gradient was followed by a 16 min re-equilibration gradient to prep the column before the next sample injection that went as follows: 0-0.05 min held at 99% B at 0.25 mL/min, 0.05-1 min from 99% B to 50% B and 0.25 mL/min to 0.1 mL/min, 1-11 min held at 50% B and 0.1 mL/min, 11-11.05 min from 50% B to 0% B at 0.1 mL/min, 11.05-14 min held at 0% B at 0.1 mL/min, 14-14.05 min held at 0% B and increased flow rate from 0.1 mL/min to 0.25 mL/min, and 14.05-16 min held at 0% B and 0.25 mL/min. The mass spectrometer acquisition settings were as follows: source voltage -2,500V, sheath gas 60, aux gas 19, sweep gas 1, ion transfer tube temperature 320°C, and vaporizer temperature 250°C. Full scan data were collected with a scan range of 70-800 m/z at a mass resolution of 240,000. Fragmentation data was collected using a data-dependent MS2 (ddMS2) acquisition method with MS1 mass resolution at 120,000, MS2 mass resolution at 15,000, and HCD collision energy fixed at 30%.

Natural isotope abundance correction was done using the FluxFix Isotopologue Analysis Tool*14* (version 0.1) web-based application^29^. Data analysis was conducted in Skyline (version 23.1.0.268), consisting of peak picking and integration, using an in-house curated compound database of accurate mass MS1 and retention time derived from analytical standards and/or MS2 spectral matches for the specified chromatography method.

### AASDHPPT co-essentiality analysis and network modeling

Gene essentiality scores and gene co-dependency Pearson correlation values were obtained from the DepMap Portal Project Achilles 23Q4 release^14,30–32^ in which genome-wide CRISPR loss-of-function screens were performed in 1100 human cancer cell lines. Co-Dependency plots between AASDHPPT and other genes were generated from gene essentiality scores for each cancer line obtained directly from the DepMap Portal. For two-dimensional network analysis, gene essentiality correlations for a custom list of genes were used to create a data matrix (see Supplementary Excel File). From this matrix, methodologies and code originally published by Arnold et al.^33^ and available at: https://github.com/finley-lab/coessentiality-network were first used to generate a correlation matrix heatmap of codependent gene modules. Co-dependent gene modules were then visualized as a network diagram and genes with low correlation scores (Pearson r < 0.2) filtered out as previously described^33^. Graph edges were weighted according to the strength of pairwise gene correlations.

### Relative quantification of NDUFAB1 species

Method was adapted from the literature for use in our model^16,17^. Standards for NDUFAB1 modifications were synthesized as in reference 14 and digested with AspN (Sigma, 11420488001) overnight at 37°C in a ratio of 1:20 (w/w) enzyme to sample, buffered to a pH of 7.6 with MOPS to a final concentration of 50mM. AspN digestion was quenched by the addition of methanol to 50%. Digested standards were then pooled for detection by liquid-chromatography mass spectrometry. For biological samples of NDUFAB1 species, ten 15-cm dishes of approximately 40% confluent C2C12 cells from each genotype were transiently transfected with either NDUFAB1-Flag or an empty vector control. After 48-hours cells were washed with ice cold PBS and frozen at -80°C. After 16 hours at -80, cells were lysed with a triton lysis buffer (1% triton, 10mM HEPES, 0.3mM EDTA, 120mM NaCl, and mammalian protease inhibitor) on ice for 30 minutes. The samples were then spun at 17,000 x g at 4°C for 30 minutes and cleared supernatants were incubated with anti-FLAG magnetic beads (Sigma, M8823) for three hours at 4°C. After three hours, bound proteins were eluted from beads using Flag peptide (Sigma, F4799) in elution buffer (10mM HEPES, 50mM NaCl, mammalian protease inhibitor) for one hour at room temperature, filtered, and quantified via Bradford assay (BioRad, 5000205). Protein that remained bound to beads was eluted with 1x Laemmli buffer and run on an SDS-PAGE gel that then underwent Coomassie staining. As determined by Bradford assay, three micrograms of Flag peptide-eluted protein from each sample were digested with AspN as above. After digest was quenched with methanol, samples were dried then resuspended in 50% methanol:50% water for further analysis by mass spectrometry. Samples were analyzed with a Vanquish (Thermo Scientific) liquid chromatography system coupled to an Orbitrap ID-X Tribrid mass spectrometer (Thermo Scientific) in full scan mode for standards and with targeted simulated ion monitoring (tSIM) for immunoprecipitation samples. For the chromatography approach, a reversed-phase C18 column (Agilent ZORBAX RRHS Extend-C18 Column, 759700-902) was used to separate samples using acidic (0.1% formic acid) solvents, and the first 1.5 minutes of column eluants were diverted to waste due to the presence of MOPS buffer in this time frame. Chromatographic traces and peak areas were quantified from extracted ion chromatograms of expected masses, matched to retention times from relevant standards using the software FreeStyle (Thermo).

### Cell immunostaining and microscopy analysis

Cells were seeded into chambered cover glass slides and cultured overnight. The following day, cells were fixed in ice-cold 4% paraformaldehyde (PFA) in PBS solution for 10 minutes at room temperature, followed by 3 washes in 1X PBS solution. Cells were then permeabilized and blocked in PBS solution containing 0.5% BSA, 5% goat serum, and 0.3% Triton X-100 for 1 hour at room temperature. Cells permeabilized/blocked with blocking buffer were then incubated with denoted primary antibodies for one hour at room temperature followed by washes with 1X PBS prior to incubation with Alexa Fluor conjugated secondary antibodies (**Table 5**) for 2 hours at room temperature. Washes were repeated and cells were then stained with Hoechst Dye for 10 minutes at room temperature. After subsequent washes, samples were then imaged on a Zeiss LSM 880 with Airyscan and non-linear optics microscope, using a 40X water objective and 1.2 NA to collect z-stacks, or with a Nikon Eclipse T*i*2 epifluorescent microscope (20X objective, 0.75 NA or 60X oil immersion objective, 1.40 NA).

For quantification of endogenous AASDHPPT expression in control and *Aasdhppt* mutant clonal lines, regions of interest (ROIs) were manually drawn with Fiji software for individual cells from channels with fluorescent ATP5a expression. Area was then measured for each ROI with corresponding mean fluorescent intensity of signal from the AASDHPPT fluorescent channel. Mean fluorescent intensity per cell area was then calculated. For statistical analysis, GraphPad Prism v10 software was used to remove outliers from the data set by the ROUT method, Q = 1%.

For colocalization analysis of endogenous AASDHPPT or AASDHPPT-GFP fusion constructs with mitochondrial proteins in maximum intensity projections of each imaged field, nuclear signal detected by Hoechst Dye staining was first used to create ROIs in Fiji. Pixels with signal detected in nuclear ROIs in red and green channels were then removed from images of the same field prior to merging of red and green channels so that colocalization analysis would be exclusive of nuclei and limited to subcellular organelles and cytoplasmic regions. Merged channels were then processed by the BIOP JACoP Fiji plugin (https://c4science.ch/w/bioimaging_and_optics_platform_biop/image-processing/imagej_tools/jacop_b/) using the Otsu method of thresholding for each channel to generate Pearsons Correlation values.

### Bimolecular fluorescence complementation (BiFC) assay

To generate BiFC cell lines, five split-Venus constructs were designed and synthesized by VectorBuilder in a pMMLV retroviral backbone. Constructs were designed to contain either the C’ terminal half of the Venus tag with blasticidin resistance (VC), or the N’ terminal half of the Venus tag with puromycin resistance (VN). VC constructs were tagged to the C’ terminal end of *citrate synthase* (CS-VC), *mitofusin 2* (MFN2-VC) or *Ndufab1* (NDUFAB1-VC). VN constructs were tagged to either the C’ or N’ terminal end of *Aasdhppt* (PPT-VN and VN-PPT, respectively). Following retroviral packaging in HEK293T cells, 40,000 wildtype C2C12 cells were transduced to express either PPT-VN or VN-PPT in combination with each VC construct. Negative control lines were generated by transducing cells with individual VC or VN constructs alongside an empty vector control (PQXCIP for VC constructs, and PQXCIB for VN constructs). This ensured that all cells were transduced with an approximately equal total viral load. Stable populations were derived via combined puromycin and blasticidin selection (2 µg/mL each), yielding 11 cell lines in total; 6 with combined VC/VN constructs plus 5 single construct controls.

For measurement of fluorescence, approximately 100,000 cells were trypsinized and resuspended in 200 µL PBS. Cell suspensions were then assessment on an Accuri bench-top cytometer, recording a minimum of 20,000 events per sample. The percentage of Venus-positive cells was analyzed using FlowJo (v. 10.10.0). Fluorescence intensity histograms were generated using FlowJo Layout Editor. A representative flow cytometry gating strategy is shown in Extended data Fig. 3C. Visual confirmation of fluorescence achieved by imaging all BiFC cell lines on a Nikon Eclipse Ti2 microscope. Briefly, 20,000 cells per cell line were seeded in chamber cover slides (Cellvis, C4-1.5H-N). After 24 hours, cells were fixed in 4% PFA for 30 mins at room temperature before imaging the following day (20X objective, 0.75 NA).

### Statistics

Statistical analysis was performed using GraphPad Prism 10. Unless otherwise noted, data were analyzed by one-way ANOVA followed by two-sided Dunnett’s multiple comparison test (when compared to only control) or Šídák’s or Tukey’s multiple comparison test (when comparing all groups). A p-value of <0.05 was considered statistically significant. Data that included a third variable (ex. time) was analyzed by two-way ANOVA followed by Tukey’s multiple comparison test.

## Abbreviations

4’PP: 4’-phosphopantetheine/yl,
AASDHPPT: aminoadipate semialdehyde dehydrogenase phosphopantetheinyl transferase,
ALDH1L2: Aldehyde Dehydrogenase 1 Family Member
L2, ATP5a: ATP synthase F1 subunit alpha
BiFC: bimolecular fluorescence complementation
CoA: coenzyme A
CS: citrate synthase
DHRS2: mitochondrial Dehydrogenase/Reductase 2
DLAT: dihydrolipoyllysine-residue acetyltransferase
DLST: dihydrolipoamide S-succinyltransferase
ETC: electron transport chain
FAS: fatty acid synthesis
FASN: fatty acid synthase
FeS: iron sulfur cluster
LA: lipoic acid
LCMS: liquid chromatography mass spectrometry
LYRM: leucine tyrosine arginine motif
mtFAS: mitochondrial fatty acid synthesis
MFN2: mitofusin 2
MTS: mitochondrial targeting sequence
NAD^+^: nicotinamide adenine dinucleotide (oxidized form)
NDUFAB1: NADH:ubiquinone oxidoreductase subunit AB1
OGDH: oxoglutarate dehydrogenase
PAGE: polyacrylamide gel electrophoresis
PDH: pyruvate dehydrogenase,
PMS: post-mitochondrial supernatant
PPT: mammalian AASDHPPT
PPTase: phosphopantetheinyl transferase
PTM: posttranslational modification
SDHB: succinate dehydrogenase subunit B
TCA: tricarboxylic acid
VC: venus c’ terminus
VN: venus n’ terminus
VUS: variant of uncertain significance

## Acknowledgments

This work was supported by funding from the NIH (R35GM151245 to SMN) and the Van Andel Institute – Metabolism & Nutrition (MeNu) Program. Stipend support for RJW and OAP was provided by the Van Andel Institute Graduate School (VAIGS). RJW was also supported by NIH F30GM154476. We thank the Meijer Foundation for financial support of MC and EM. We also thank the Van Andel Institute’s Flow Cytometry Core (RRID:SCR_022685) for their assistance with single cell sorting for generation of clonal cell lines, the Metabolomics and Mass Spectrometry Core (RRID:SCR_024903) for their assistance with labeled glutamine tracing experiments and analysis as well as method development for the direct relative quantification of NDUFAB1 acylation species, the Optical Imaging Core (RRID:SCR_021968) for their assistance with airyscan microscopy, and the Bioinformatics and Biostatistics Core (RRID:SCR_024762) for their assistance with gene essentiality score correlation network analysis.

## Author Contributions

Conceptualization: PRN, RJW, SMN; Methodology: PRN, RJW, MC; Investigation: PRN, RJW, SEJP, MC, NG, JZL, OAP, EM; Formal Analysis: PRN, RJW, SEJP; Funding acquisition: RJW, SMN; Resources: SMN; Supervision: SMN; Visualization: PRN, RJW, SEJP; Writing – original draft: PRN, RJW, JZL, SMN; Writing – review & editing: PRN, RJW, SEJP, MC, OAP, MLH, SMN

The authors declare no competing interests.

## Data Availability

All data needed to evaluate the conclusions in the manuscript are present in the manuscript and/or Supplementary Information. Additional data related to this paper may be requested from the authors.

---

**Extended Data Fig. 1.**
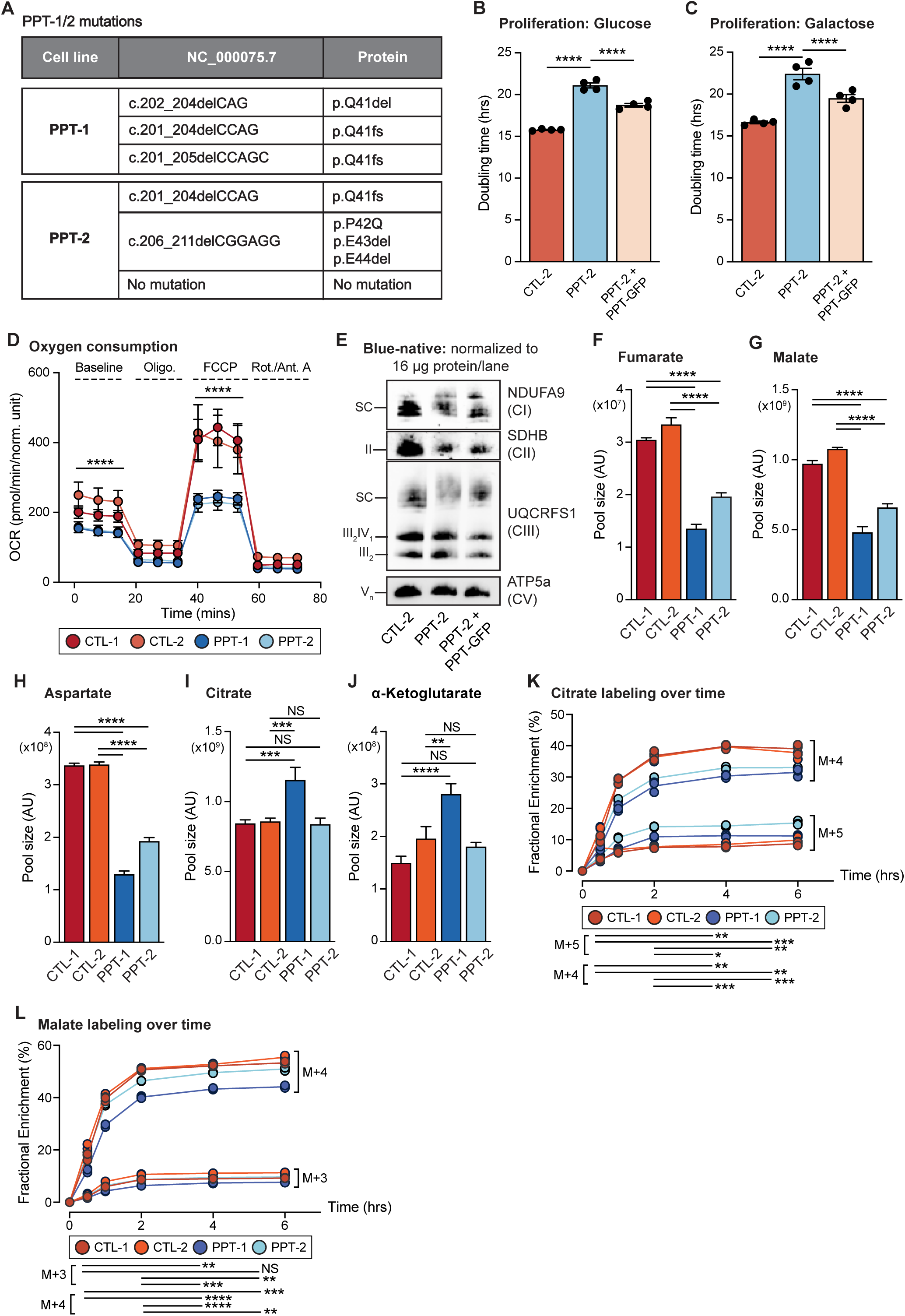
AASDHPPT is required for mitochondrial oxidative metabolism. ((A) Sequence analysis sgRNA-targeted regions PCR-amplified and subcloned into pCR™4-TOPO plasmid in clonal *Aasdhppt* mutant cell lines, and corresponding mutations sequenced. (B-C) Cells were grown in either 25 mM glucose (B) or 10 mM galactose (C) in an IncuCyte® system with images captured every 3 hours for 4 or 2 days respectively. Data represent the mean doubling time calculated from technical replicates of growth curves; error bars denote SEM, **** = p <0.0001 as determined by one-way ANOVA with Tukey’s multiple comparisons test. Data are representative of 1 experiment from n > 3 biological replicates. (D) Seahorse mitochondrial stress test for Oxygen Consumption Rate (OCR) of indicated cell lines. Data are representative of 1 experiment from 3 biological replicates. Error bars represent SD. Significance determined at indicated time points using one-way ANOVA, **** = p <0.0001 (E) Blue-native PAGE separation of isolated mitochondrial protein complexes from denoted cell lines, followed by immunoblot with the indicated antibodies. Data are representative of n > 3 biological replicates. (F-J) Pool size of the indicated metabolites from steady state metabolomics analysis at the four-hour labeling time point. Data from n=3 technical replicates of the indicated cell lines. Error bars represent SD. Statistical analysis was done using a one-way ANOVA followed by Tukey’s multiple comparisons test. (K-L) Fractional enrichment of given isotopologues is represented as percentage of the total pool at each indicated time point. Error bars represent SD. Significance was determined by two-way ANOVA followed by Tukey’s multiple comparisons test. ** = p <0.01. *** = p <0.001. **** = p <0.0001. NS = not significant.

**Extended Data Fig. 2.**
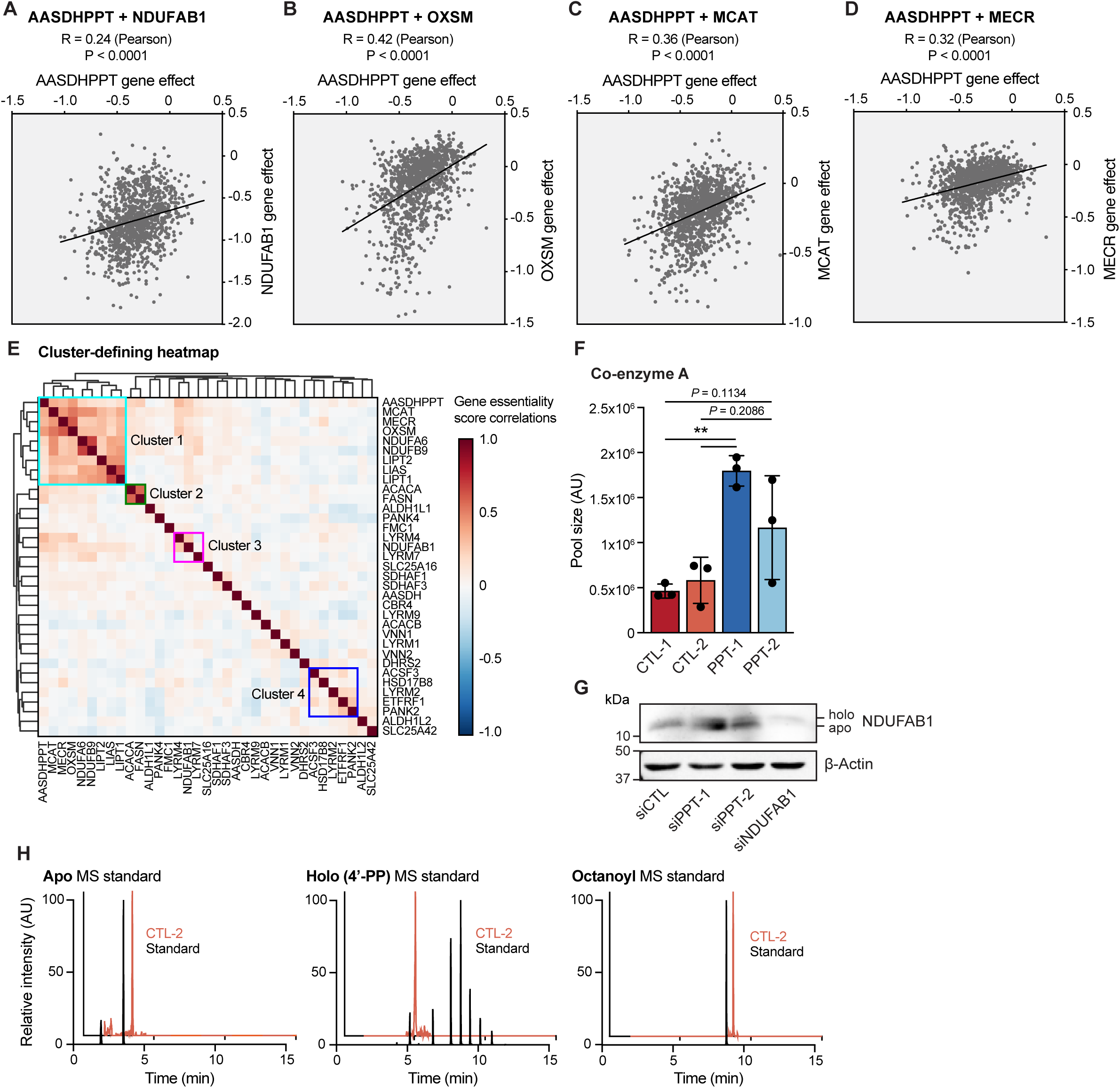
AASDHPPT is required for mitochondrial fatty acid synthesis pathway function. (A-D) Codependency plots of gene essentiality scores comparing *AASDHPPT* with *NDUFAB1* (A), *OXSM* (B), *MCAT* (C), and *MECR* (D) from DepMap CRISPR screens^14,30–32,34^**. (**E) Heatmap depicting hierarchical clustering of pairwise gene essentiality score correlations of PPT, known PPT substrate targets, mtFAS pathway members, LYRM family members, and genes associated with the GO term “pantothenate metabolism.” (F) CoA pool size in cells of the indicated genotype. Statistical analysis performed via one-way ANOVA with Tukey’s multiple comparisons. ** = p <0.01, other p values as written. (G) Whole cell lysates from WT cells transfected with siRNAs targeting *Aasdhppt, Ndufab1*, or scramble control were separated by SDS-PAGE and immunoblotted for the denoted targets. Data are representative of 3 biological replicates. H) Representative extracted ion chromatograms for indicated NDUFAB1 species from control cell line overlayed with extracted ion chromatograms from a pool of Apo-Holo- and Acyl-standards run with identical chromatography. Standards were run in full scan mode while samples were run with targeted selective ion monitoring.

**Extended Data Fig. 3.**
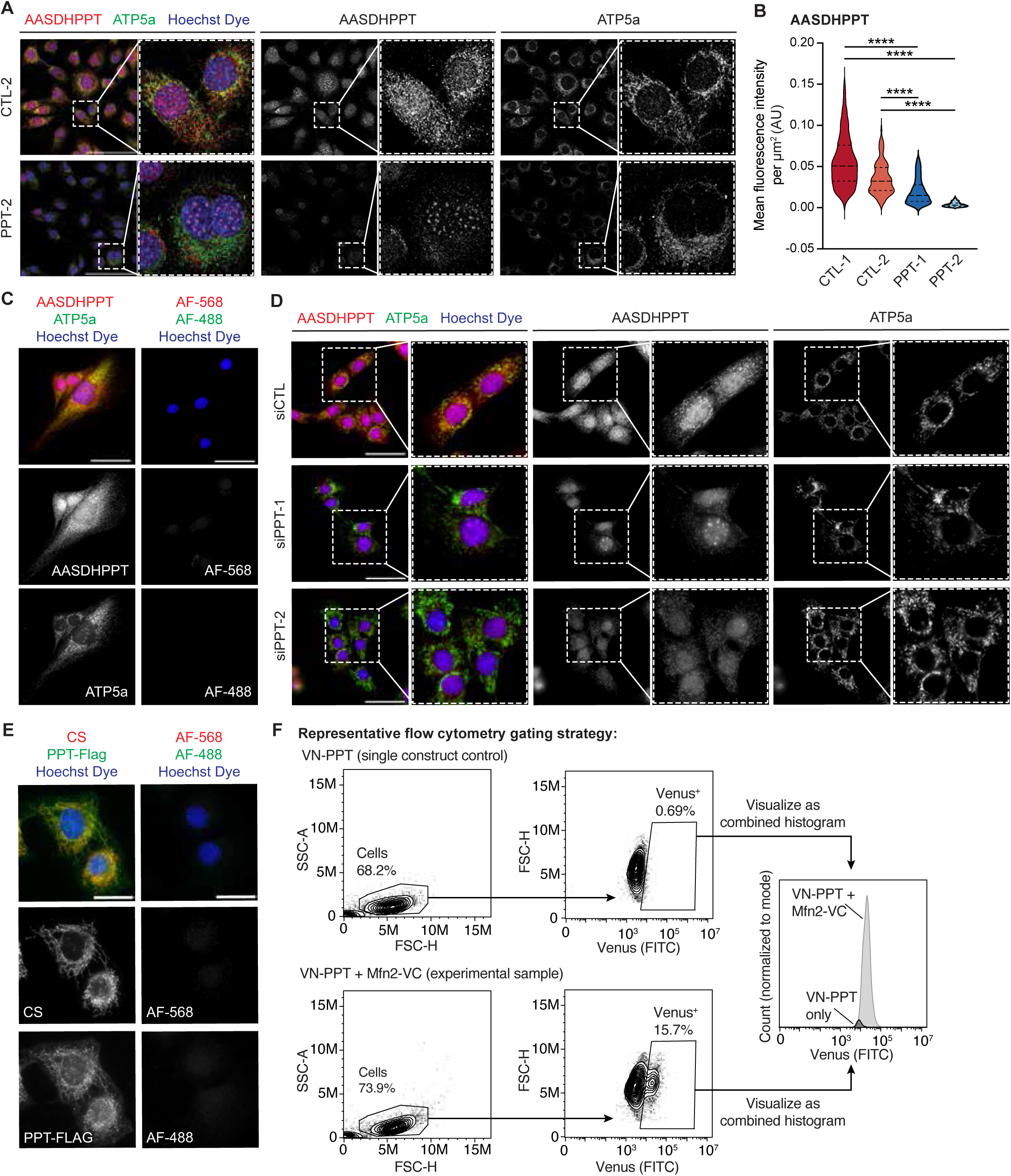
AASDHPPT localizes to the mitochondrial matrix. (A) Representative 40X HPF images of the indicated cell lines stained with Hoechst dye and antibodies against AASDHPPT (red) and ATP5a (green). Scale bars = 100µm. (B) Quantification of mean fluorescent intensity of AASDHPPT expression per cell area (μm^2^) in *Aasdhppt* mutants (PPT-1 n = 170 and PPT-2 n =159) and control clonal cell lines (Ctrl-1 n = 186 and Ctrl-2 n = 118) as represented in A and Fig. 3A. (C) Representative 60X HPF images of WT cells stained with Hoechst dye and antibodies against AASDHPPT (red) and ATP5a (green) along with secondary antibody-only controls. Scale bars are 25 μm. (D) Representative 20X HPF images of WT cells transfected with the denoted siRNAs following fixation at 72 hours post-transfection. Cells were stained with Hoechst dye and antibodies against AASDHPPT (red) and ATP5a (green). Scale bars are 50 μm. (E) Representative 60X HPF images of WT cells expressing AASDHPPT-Flag (PPT-Flag) stained with Hoechst dye and antibodies against Citrate Synthase (red) and Flag-tag (green) along with secondary antibody-only controls (AF-568 and AF-488). Scale bars are 25 μm. (F) Representative schematic of flow cytometry gating strategy for experiments in Fig. 3E-F.

**Extended Data Fig. 4.**
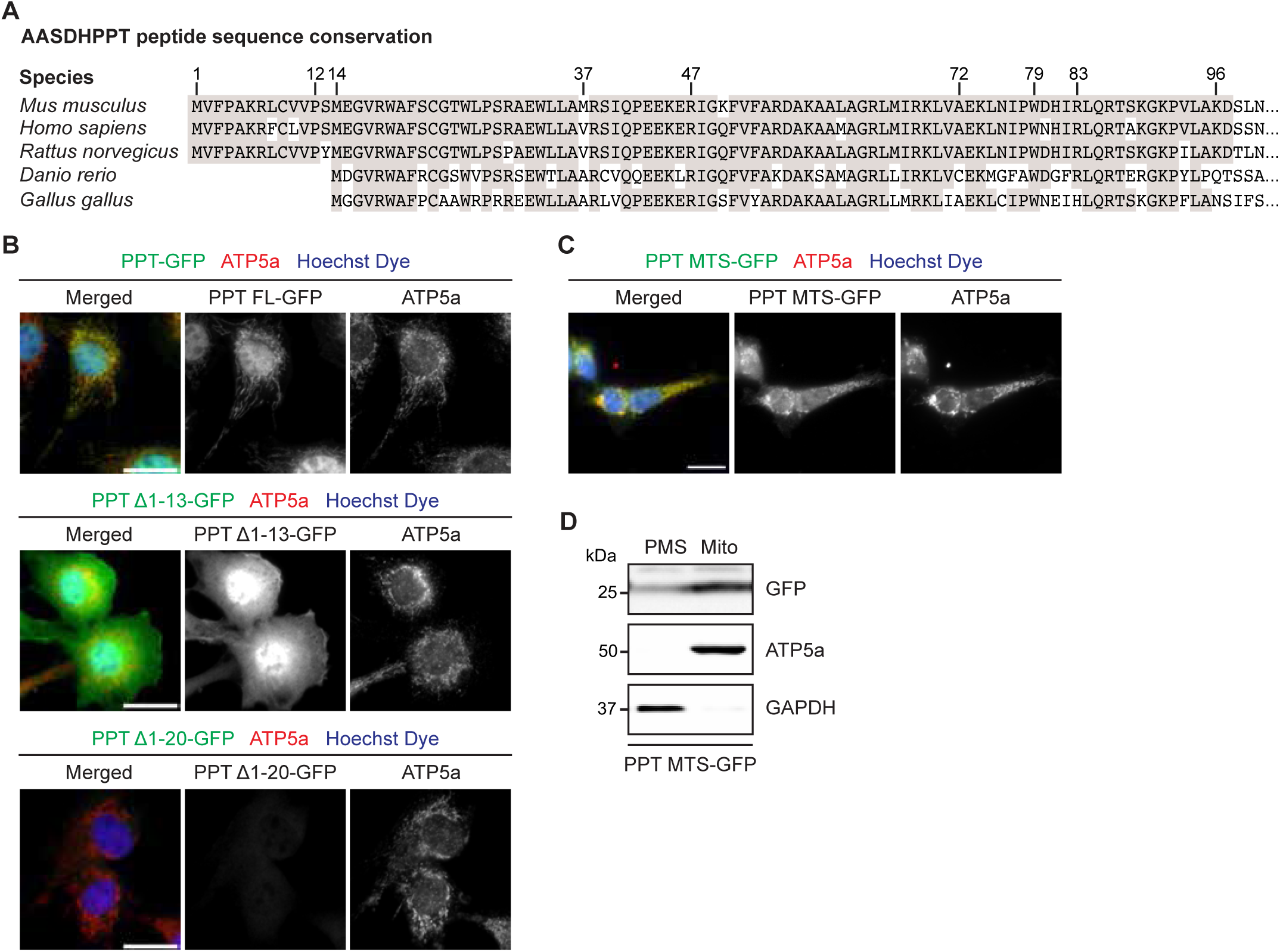
AASDHPPT has a N-terminal MTS. (A) Sequence alignment: MTS conservation. Accession numbers as follows; *Mus musculus:* NP_001313288.1, *Homo sapiens: NP_056238.2, Rattus norvegicus: NP_001100268.2, Danio rerio: NP_001028901.1, Gallus gallus: NP_001265017.1*. (B) Representative 60X HPF images of WT cells expressing PPT-GFP, 13 residue N-terminal truncation of PPT (PPT_∆1-13_-GFP), or 20 residue N-terminal truncation of PPT (PPT_∆1-20_-GFP) constructs, stained with Hoechst dye and antibody against ATP5a (red). Scale bars are 25 μm. (C) Representative 60X HPF images of cells expressing PPT MTS-GFP stained with Hoechst dye and antibody against ATP5a (red). Scale bars represent 20 μm. (D) Subcellular fractionation of cells expressing PPT MTS-GFP. Post-mitochondrial supernatant (PMS) and mitochondrial lysate (Mito) was isolated, separated by SDS-PAGE, and immunoblotted for the indicated targets. Blots are representative of 3 biological replicates.

**Extended Data Fig. 5.**
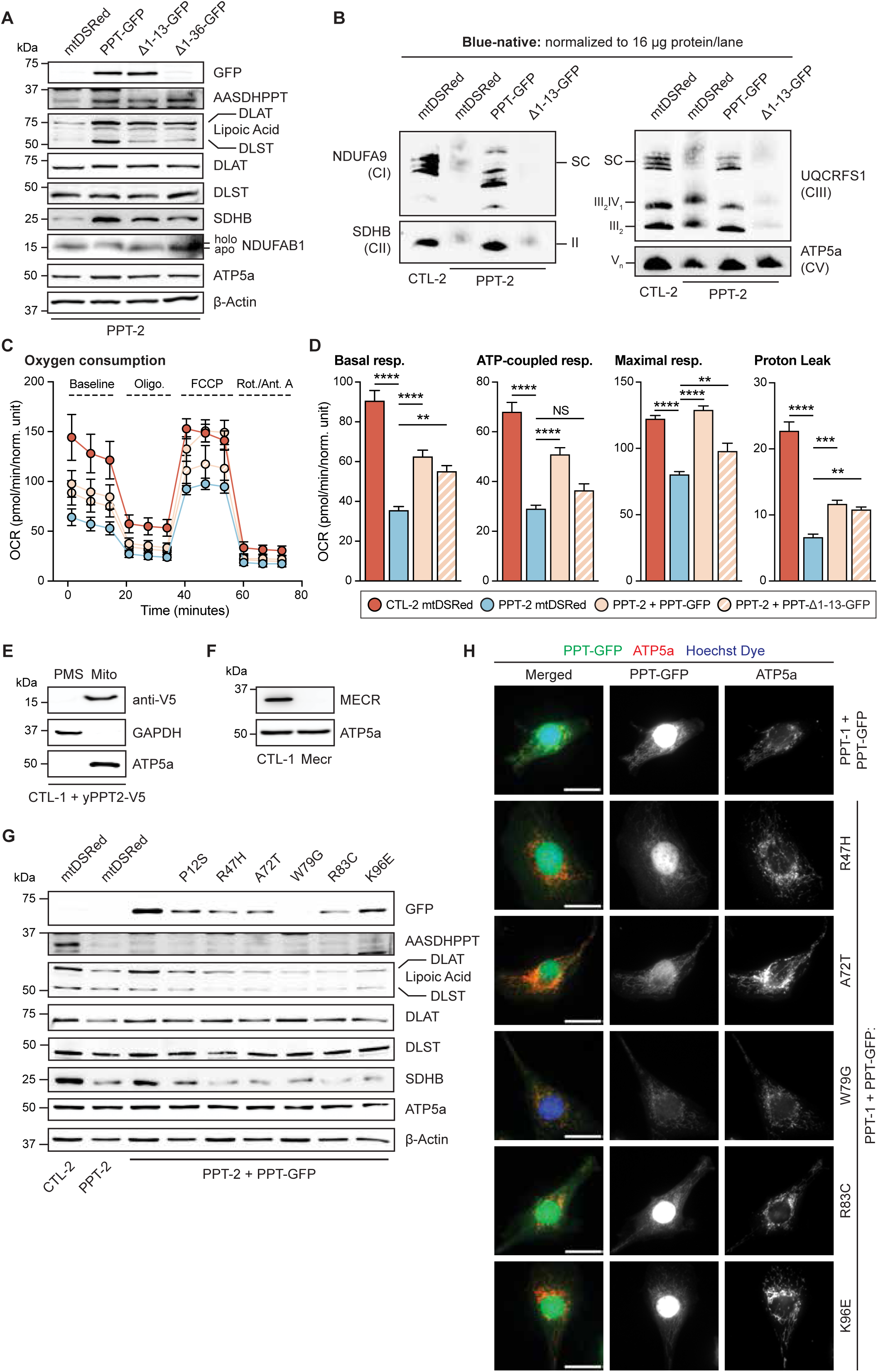
Mitochondrial localization of AASDHPPT is required for mtFAS function. (A) Whole cell lysates from cells expressing mtDSRed, PPT-GFP, PPT_∆1-13_-GFP, or PPT_∆1-36_-GFP were separated by SDS-PAGE and immunoblotted for the denoted targets. Data are representative of n > 3 biological replicates. (B) Blue-native PAGE separation of isolated mitochondrial protein complexes from denoted cell lines expressing mtDSRed, PPT-GFP, or PPT_∆1-13_-GFP, followed by immunoblot with the indicated antibodies. Data are representative of n > 3 biological replicates. (C) Seahorse mitochondrial stress test for Oxygen Consumption Rate (OCR) of indicated cell lines expressing mtDSRed, PPT-GFP, or PPT_∆1-13_-GFP. Data are representative of 1 experiment from 3 biological replicates. Error bars represent SD. (D) Quantification of basal respiration, ATP-production coupled respiration, maximal respiration and proton leak from data in (C). Data are mean OCR; error bars are SEM. * = p <0.05, ** = p < 0.01, *** = p < 0.001, **** = p <0.0001, determined by one-way ANOVA with Tukey’s multiple comparisons test. (E) Subcellular fractionation of clonal control cells expressing yPpt2-V5. Post-mitochondrial supernatant (PMS) and mitochondrial lysate (Mito) was isolated, separated by SDS-PAGE, and immunoblotted for the indicated targets. Data are representative of 3 biological replicates. (F) Whole cell lysates from clonal control *and* Mecr-deficient cells were separated by SDS-PAGE and immunoblotted for the denoted targets. Data are representative of 3 biological replicates. (G) Whole cell lysates from indicated cell lines expressing mtDSRed, PPT-GFP P12S, PPT-GFP R47H, PPT-GFP A72T, PPT-GFP W79G, PPT-GFP R83C, or PPT-GFP K96E were separated by SDS-PAGE and immunoblotted for the denoted targets. Immunoblots are representative of n>3 biological replicates. (H) Representative 0X HPF images of WT cells expressing PPT-GFP, PPT-GFP R47H, PPT-GFP A72T, PPT-GFP W79G, PPT-GFP R83C, or PPT-GFP K96E stained with Hoechst Dye and antibody against ATP5a (red). Scale bars represent 25 μm.

